# Metastasis-associated wound repair promotes reciprocal activation of the lung epithelium and breast cancer metastases during outgrowth

**DOI:** 10.1101/2025.07.23.666437

**Authors:** Jessica L. Christenson, Nicole S. Spoelstra, Michelle M. Williams, Kathleen I. O’Neill, David J. Orlicky, Jennifer A. Wagner, Andrew E. Goodspeed, Li Wei Kuo, Lyndsey S. Crump, Jennifer K. Richer

## Abstract

When tumor cells colonize distant organs during metastasis, they interact extensively with surrounding cells. These interactions often change the behavior of surrounding cell populations which collectively induce a pro-tumor microenvironment that permits tumor cell outgrowth into overt, clinically detectable metastatic disease. The lung is one of the most common sites of breast cancer (BC) metastasis. A chronic wound repair-related phenotype developed within the lung microenvironment during metastatic outgrowth in immunocompetent preclinical mouse models of BC. This phenotype was characterized by an increased number and activation of lung type II alveolar epithelial (AT2) cells surrounding growing metastases. Metastatic outgrowth significantly changed AT2 gene expression, resulting in a modified secretome. AT2-derived secreted factors also promote TNBC growth. AT2 secreted factors are regulated by the cAMP response element-binding protein (CREB). Targeting CREB signaling with the phosphodiesterase 4 (PDE4) inhibitor roflumilast reduced AT2-BC reciprocal interactions in vitro and metastatic outgrowth in vivo.

**STATEMENT OF SIGNIFICANCE:** Alveolar epithelial cells are the most common cell type in the lung. Our studies demonstrate the potential for targeting metastasis-associated wound repair and lung epithelial cell activation during metastatic outgrowth with FDA-approved PDE4 inhibitors. This strategy may be an effective way to treat and manage progression of established metastatic BC.

## INTRODUCTION

Metastatic outgrowth is one of the most clinically relevant stages of the metastatic cascade [1]. During this stage, metastatic disease becomes radiologically detectable, patients are diagnosed, and treatment begins. While therapeutic interventions used to treat metastatic disease are often based on characterization of a patient’s primary tumor, since many metastases are difficult to biopsy, there is a significant amount of research indicating that metastases are distinct and may require specifically tailored therapies [2, 3]. Thus, it is critical that treatment strategies be developed to specifically target this phase of metastatic progression to reduce the mortality associated with metastases by: 1) preventing destructive metastatic outgrowth, 2) blocking secondary metastatic spread, and 3) improving metastatic organ function. To achieve this goal, it is imperative that we investigate how the metastatic microenvironment is altered during disease progression and how this supports metastatic outgrowth.

For patients with breast cancer (BC), tumors primarily metastasize to the bone, lung, liver and brain [4], and the median overall survival for patients with metastatic BC is 1-4 years, depending on subtype [5, 6]. Approximately one third of people with metastatic BC develop lung metastases [7]. Aggressive BC, such as triple-negative BC (TNBC), that metastasize within the first few years post-diagnosis of primary disease, preferentially metastasize to the lung [8], while other subtypes commonly exhibit secondary metastatic spread to the lung from sites like the bone [9]. Metastatic BC cells colonize alveoli, the terminal structures of the lung [10] where they cause injury to the alveolus and vulnerable type I alveolar epithelial (AT1) cells as they grow. The specific contributions of this damage to metastatic progression, however, remain unknown. AT1 cells are responsible for gas exchange and are critical for proper lung function. AT1 cell damage triggers the release of secreted factors that rapidly initiate a wound healing cascade. This cascade is characterized by a distinct series of events, including inflammation caused by the accumulation of neutrophils and macrophages, the recruitment of fibroblasts that deposit collagen, and the proliferation of lung type II alveolar epithelial (AT2) cells that terminally differentiate into AT1 cells to repopulate the wound [11].

AT2 cells are often called the “defenders of the alveolus” and have three primary functions, which include: 1) facilitating AT1 gas exchange through the secretion of surfactant to reduce surface tension within the alveolar airspace to prevent lung collapse during breathing, 2) acting as the master regulators of injury and repair within the lung by initiating and coordinating the repair process, and 3) serving as the stem cells of the alveolar epithelium, a necessary component of wound resolution following injury [12]. Critical immunological coordination by AT2 cells is achieved through interactions, both direct or indirect, with innate and adaptive immune cells. AT2 cells have the capacity to recruit neutrophils, activate macrophages, and even present antigen to T cells (as reviewed in [1]). AT2 cells are the most numerous cells within the alveolus [13] and are likely to interact extensively with metastatic cells in the lung. Some studies suggest that AT2 cells may play a role in cancer progression [14–16], but none have investigated AT2 behavior in the context of BC metastatic outgrowth. We hypothesized that BC lung micrometastases activate surrounding lung epithelial cells which, in turn, support the outgrowth of metastases.

Our findings demonstrate that chronic wound repair develops during metastatic outgrowth, and that growing metastases alter AT2 cell function in metastasis-adjacent tissue. Activated AT2 cells, in turn, promote metastatic outgrowth through secreted signaling peptides. Most importantly, we were able to impede BC-AT2 reciprocal activation, and consequent metastatic outgrowth, by targeting pro-inflammatory phosphodiesterase 4 (PDE4) activity using the FDA-approved inhibitor roflumilast (ROF), a drug currently used to treat patients with chronic obstructive pulmonary disease (COPD) (Fig. 1).

**Figure 1.**
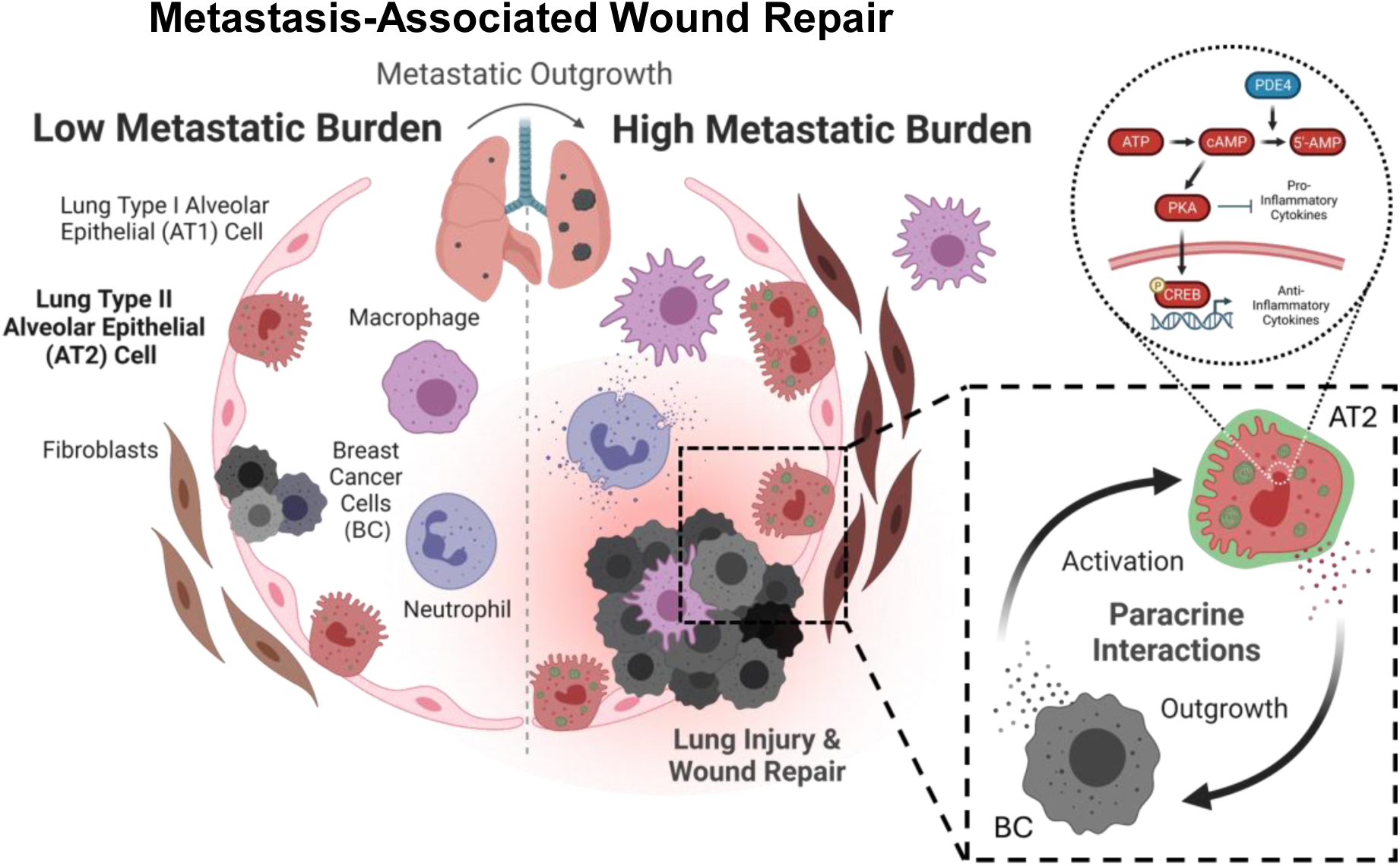
Metastasis-associated wound repair in the lung. Summary model of study results. During metastatic outgrowth, breast cancer (BC) micrometastases grow within the lung alveoli and cause injury to surrounding type I alveolar epithelial (AT1) cells. Chronic wound repair develops throughout this process which is characterized by an increase in the number and activation of adjacent resident lung cells, including epithelial, immune and stromal cells. Type II alveolar epithelial (AT2) cells seem to play a prominent role in metastatic outgrowth as a known regulator of wound repair within the lung. During outgrowth, AT2 cells reciprocally interact with adjacent BC cells through paracrine mechanisms. Several AT2 secreted factor genes are known cAMP response element-binding protein (CREB)-regulated genes. Phosphodiesterase 4 (PDE4) inhibition may be a rational and effective strategy for blocking AT2-BC interactions and metastatic outgrowth. cAMP, cyclic adenosine monophosphate; ATP, adenosine triphosphate; AMP, adenosine monophosphate; PKA, protein kinase A. Created with BioRender.com.

## MATERIALS AND METHODS

### Reagents

Roflumilast (ROF) was purchased from Selleck Chemical LLC. (No. S2131), and cilomilast (CILO) was purchased from MedChemExpress (No. HY-10790). For in vitro experiments, ROF and CILO were diluted in dimethyl sulfoxide (DMSO). For in vivo experiments, ROF was diluted in a vehicle consisting of 30% polyethylene glycol 400 (PEG400), 0.5% Tween 80, and 5% propylene glycol prepared in sterile deionized water (diH_2_O).

### Cell culture

All cells were cultured in 5% CO_2_. SUM159PT and MDA-MB-453 (MDA453) TNBC cells were purchased from the University of Colorado Cancer Center (UCCC) Cell Technologies Shared Resource in 2008 and the American Type Culture Collection (ATTC) in 2012, respectively. Both cell lines were maintained in DMEM with 10% fetal bovine serum (FBS). BT549 TNBC cells were purchased in 2008 from ATCC and maintained in RPMI 1640 with 10% FBS. Aside from A549 cells, all cells were passaged/subcultured <10 times post-thaw. A549 lung carcinoma cells were purchased in 2019 from the UCCC Cell Technologies Shared Resource and maintained in F-12K media with 10% FBS. A549 cells were aged to a more AT2-like phenotype by culturing for a minimum of 8 passages [17]. Human induced pluripotent stem cells (iPSC), originating from male foreskin fibroblasts, were purchased from the Gates Institute at the University of Colorado Anschutz Medical Campus (CU Anschutz) in 2019. iPSC were maintained on matrigel-coated plates in mTeSR1 media (Stemcell Technologies, No. 85850) for a maximum of 5 passages. iPSC cells were differentiated into AT2 cells (iAT2) following the protocol outlined in Tamò et al. 2018 [18]. Briefly, iPSC cells were plated on vitronectin XF (Stemcell Technologies, No. 07180)-coated plates for 3 days in mTerS1 media. iPSC were differentiated into endoderm cells by culturing in STEMdiff definitive endoderm media (Stemcell Technologies, No. 05110) for 4 days. Endoderm cells were then trypsinized, counted, and plated for experimentation in SAGM media (small airway epithelial cell growth medium; Lonza Walkersville, No. CC-3118) supplemented with 1% FBS for 3 days to induce differentiation into lung AT2 cells. iAT2 cells were cultured for no more than 21 days to avoid transdifferentiation to AT1 cells. Met-1 mouse mammary carcinoma cells were kindly provided in 2015 by Donald McDonnell (Duke University) with permission granted by Alexander Borowsky (University of California – Davis). Met-1 cells were maintained in DMEM with 10% FBS. All cell lines were routinely tested for mycoplasma contamination and authenticated in 2025 by short tandem repeat DNA profiling by the UCCC Cell Technologies Shared Resource. Transwell plates with 0.4µm pores were used to investigate paracrine interactions between cell lines (no contact co-culture), and AT2 media was used for all co-culture experiments. Cell numbers and timeframes for each experiment, depending on cell type and assay, can be found in Supplementary Table 1.

#### Cell viability

The crystal violet cell viability assay was used to examine how no contact co-culture or ROF treatment affects cell numbers. Cells were fixed with 10% buffered formalin and stained with 0.1% crystal violet dye prepared in 25% methanol. Stained cells were scanned for analysis of confluence using ImageJ software (NIH) to calculate the percent area per well covered by cells. Crystal violet dye was then solubilized with 10% acetic acid and absorbance was measured at 570nm to quantify relative cell number. Data was presented as relative cell number normalized to the mean confluence or mean absorbance of cells cultured alone (for co-culture experiments) or DMSO-treated cells (for ROF experiments).

#### Lysotracker stain

For fluorescent imaging, cells were plated in 6-well plates and for fluorescent quantification experiments, cells were plated in 24-well plates. In live cells, nuclei were stained with 2µg/mL Hoechst for 30min at 37°C. 50µM LysoTracker Green DND-26 (Cell Signaling Technology, No. 8783), prepared in Opti-Klear Live Cell Imaging Buffer (abcam, No. ab275939), was then used to stain AT2 lamellar bodies. Representative images were taken using a BX40 microscope, DP73 camera, and cellSens standard software (Olympus). Alternatively, fluorescence was measured at an excitation/emission of 485/524nm for lysotracker and 360/460nm for Hoechst. Relative to cells cultured alone, co-cultured cell data was presented as mean lysotracker fluorescence normalized to Hoechst fluorescence.

#### Cell morphology

Cells were grown on glass coverslips in 6-well plates. Coverslips were sterilized with a wash of 70% ethanol (EtOH), air dried for 10min, and UV exposed for 10min. A549 cells were grown on uncoated coverslips. iAT2 cells were grown on poly-L-lysine (Millipore Sigma, No. P8920)-coated coverslips. At the experimental endpoint, cells were rinsed in 1X phosphate buffered saline (PBS), fixed in 10% formalin for 5min and 50% EtOH for 4min, and washed in 1X PBS for 5min. Cells were hematoxylin and eosin (H&E) stained using an abbreviated protocol. Briefly, coverslips were washed in 100% EtOH for 5min, stained in 1% Eosin-Y (Cancer Diagnostics Inc., No.832) for 1min, washed twice with tap water, stained in Harris Hematoxylin (Cancer Diagnostics Inc., No.842) for 5min, washed with tap water, and mounted on a slide using Permount mounting medium (Fisher Scientific, No. SP15-100).

### In vivo mouse models

All in vivo experiments were performed in accordance with NIH Guidelines of Care and Use of Laboratory Animals. Mice were euthanized by CO_2_ inhalation followed by cervical dislocation. Tissues were fixed overnight with 10% buffered formalin for histological analysis. Lungs were intratracheally inflated with formalin prior to dissection to maintain the structural integrity of the lung architecture. Fixed tissues, stored in 70% ethanol, were then processed and paraffin-embedded by the UCCC Pathology Shared Resource (PSR).

#### Transgenic spontaneous metastasis model

Formalin-fixed, paraffin embedded (FFPE) MMTV-PyMT (mammary specific polyomavirus middle T antigen) metastatic lung tissue was kindly provided by Susan Kane (City of Hope) in 2014. Tissue was collected from 20-week-old hemizygous, mixed background (C57BL/6 and FVB) MMTV-PyMT females with multiple mammary tumors.

#### Late-stage metastasis model

To study the latter stages of lung metastasis, such as metastatic outgrowth, 5×10^5^ Met-1 mouse mammary carcinoma cells in 100µL 1X PBS were intravenously (IV) injected into the tail veins of 10-week-old female FVB/NJ mice (The Jackson Laboratory, No. 001800). Alternatively, 3×10^5^ 66Cl4 mouse mammary carcinoma cells in 100uL 1X PBS were IV injected into the tail veins of 10-week-old female BALB/cJ mice (The Jackson Laboratory, No. 000651). To examine how the metastatic microenvironment is altered during outgrowth, lungs with a low versus high metastatic burden were compared. Lungs were collected 1 week following IV injection as a model for low metastatic burden lungs and 3 weeks post-injection as a model for high metastatic burden. The effects of ROF on metastatic outgrowth were examined using a late-stage metastasis model (n=5-8 mice/group). Three days post-injection of Met-1 cells, mice began daily oral treatment of ROF (5mg/kg/day) or vehicle control for 3 weeks. Mouse weights were measured weekly.

#### Lung magnetic resonance imaging (MRI)

To confirm metastatic burden prior to tissue homogenization for single cell RNA-sequencing (scRNAseq), the lungs of mice were MRI scanned by the UCCC Animal Imaging Shared Resource (AISR). Mice underwent high-resolution breath- and cardiac-gated MRI [19]. MRI was performed on anesthetized mice (1.5% isoflurane) using a 9.4 Tesla BioSpec animal MRI scanner (Bruker Medical).

### Histology

5µm thick sections of formalin-fixed paraffin-embedded (FFPE) tissue samples were used for analyses. Standard procedures were followed for H&E staining using Harris Hematoxylin and Eosin-Y. Neutrophils and macrophages surrounding small versus large metastases were histologically counted by an experienced veterinary pathologist (Linda Kassenbaum Johnson, University of Colorado).

#### Immunohistochemistry (IHC)

Sections were deparaffinized in a series of xylenes and ethanols. 1X tris-buffered saline (TBS) with 0.05% Tween 20 was used for all washes. Each antibody was optimized to determine the best retrieval buffer, antibody dilution, and detection method. See Supplementary Table 1 for the IHC protocol details for each antibody used. Antigen heat retrieval was performed using one of three buffers, depending on the antibody: 1) 10mM citrate buffer, pH 6.0; 2) 10mM Tris/1mM ethylenediaminetetraacetic acid (EDTA), pH 9.0; or 3) 1X Dako target retrieval solution (TRS) citrate buffer, pH 6.0 (Agilent Technologies, No. S169984). Antibodies were detected using one of four methods: 1) Vector ImmPRESS anti-rat polymer HRP (Vector Laboratories, No. MP-7444); 2) Vector ImmPRESS anti-rabbit polymer HRP (Vector Laboratories, No. 7451); 3) Dako REAL EnVision polymer HRP rabbit-mouse (Agilent Technologies, No. K4003); and 4) Jackson biotinylated goat anti-rat IgG (Jackson ImmunoResearch Laboratoies Inc., No. 112-065-003) followed by streptavidin HRP. The ImmPACT DAB peroxidase (HRP) substrate kit (Vector Laboratories, No. ZJ0415) was used for detection, and cells were counterstained with a 1:5 dilution of Harris Hematoxylin. Slides were mounted using Permount mounting medium. Representative images were taken using a BX40 Microscope (Olympus) with a SPOT Insight Mosaic 4.2 camera and software (Diagnostic Instruments, Inc.).

#### Metastatic burden quantification

To get an accurate representation of metastatic burden following treatment, mouse lungs were serial sectioned and three sections, separated by 50µm, were stained for PyMT by IHC. Whole slides were imaged using the Aperio Digital Pathology Slide Scanner (Leica Biosystems) by the UCCC PSR. The number and size (area in µm^2^) of metastases were quantified per slide using the Aperio ImageScope software (Leica Biosystems). Data is presented as the average per mouse calculated from all three sections.

#### Picrosirius red staining

Tissues were stained by the UCCC PSR using a standardized protocol. Briefly, slides were deparaffinized in a series of xylenes and ethanols. Slides were stained with Weigert’s Hematoxylin for 8min, washed in tap water for 10min, stained in picrosirius red solution for 1hr, rinsed with acidified water, and coverslipped with mounting medium. Representative images were taken under white light and polarized light using an Olympus BX51 microscope equipped with a four-megapixel Macrofire digital camera (Optronics) using the PictureFrame Application 2.3 (Optronics).

#### Multispectral immunofluorescence (multi-IF)

Sections were deparaffinized in a series of xylenes and ethanols. 1X TBS with 0.05% Tween 20 and diH_2_O were used for washes. See Supplementary Table 1 for protocol and antibody details. Primary antibodies were detected with Opal TSA technology (Akoya Biosciences). Slides were scanned using a Vectra Polaris Quantitative Pathology Imaging System (Akoya Biosciences). Regions of interest (ROI) were selected for analysis at the periphery of lung metastases to focus on the adjacent microenvironment. Multispectral images were unmixed, and tissue/cellular compartments were segmented before cells were assigned phenotypes using inForm Tissue Analysis software (Akoya Biosciences).

### Cytokine array

Using the late-stage metastasis model with Met-1 cells, lungs were collected from mice with a low versus high metastatic burden and snap frozen in liquid nitrogen (n=3 mice/group). The left lobes from each lung were homogenized and protein was isolated for cytokine analysis using the Proteome Profiler Mouse Cytokine Array Kit (R&D Systems, No. ARY006) according to the manufacturer’s instructions. Protein expression was quantified using ImageJ software and expressed as mean pixel density per mouse.

### Quantitative PCR (qPCR)

QIAshredder columns (Qiagen, No. 79654) were used to prepare cell lysates and the RNeasy Plus Kit (Qiagen, No. 74136) was used to isolate RNA. cDNA was synthesized using iScript Reverse Transcription Supermix (Bio-Rad, No. 1708840). SYBR Green quantitative gene expression analysis was performed using the PowerUp SYBR Green Master Mix (ThermoFisher Scientific, No. A45742) on the 7500 Fast Real-Time PCR System (Applied Biosystems). See Supplementary Table 1 for primer details. Relative gene expression was calculated using the comparative cycle threshold method and values were normalized to *18S*.

### Bioinformatics

#### scRNAseq

Using the late-stage metastasis model with Met-1 cells, lungs were collected from mice with a low versus high metastatic burden (n=1 mouse/group). Mouse lungs were enzymatically dissociated. Approximately 10,000 cells were used to generate libraries and sequenced by the Genomics Core facility at the University of Colorado Anschutz Medical Campus. FASTQ files for each sample were processed using Cell Ranger 2.0.2. (10x Genomics) using a modified version (PYMT was appended) of the mm10 reference genome. The resulting data was then aggregated using the Cell Ranger aggr pipeline. A total of 5,765 cells (high-burden n=3,121; low-burden n=2,644) remained following aggregation. Expression data for high-burden and low-burden cells was imported into an R environment; the data was filtered and analyzed using the R packages Seurat and Monocle. Filtering based on unique gene counts less than 1,000 or greater than 7,000, or mitochondrial percentages greater than 10% resulted in 5,504 cells in the final analysis, with an average of 11,764 UMI counts per cell and a median of 2,487 genes per cell. Principal Component Analysis (PCA) was then performed, and the top 10 principal components (PC) were selected for subsequent use. The dimensionality was reduced to two dimensions using t-stochastic neighbor embedding (t-SNE), with the 10 PCs being used as input. Cell clusters were demarcated via fast search and find of density peaks [20]. Differentially expressed genes were identified for each cluster using the two-sided Student’s *t*-test provided by the Seurat FindMarkers function. Normalized gene expression within cells from a particular cluster was compared to expression within cells from all other clusters. Genes with less than a 25% change or those expressed in less than 10 percent of cells in both populations being compared were excluded from analysis. Following cluster designation, a likelihood ratio test using a generalized linear model was performed to identify genes that vary by cluster. Genes with an adjusted *p*-value of 0.05, an average greater than the bottom quintile of averages, and a dispersion higher than what would be expected using the DESeq model were then selected to construct a developmental trajectory using DDRTree. These data are available in the Gene Expression Omnibus (GEO) database as [*in progress*]. Transcriptomics data were then compared to a published tissue dissociation-related gene signature to rule out enzymatic dissociation as a major contributor to gene expression changes between groups [21].

#### Bulk RNAseq

iAT2 cells were no contact co-cultured with TNBC cells for 5 days. QIAshredder columns (Qiagen, No. 79654) were used to prepare cell lysates and the RNeasy Plus Kit (Qiagen, No. 74136) was used to isolate RNA. Illumina HiSeq libraries were prepared and sequenced by the Genomics and Microarray Core Facility at CU Anschutz and the UCCC Bioinformatics and Biostatistics Shared Resource assisted with data analysis. Briefly, Illumina adapters and the first 12 base pairs of each read were trimmed using BBDuk (BBMAP; www.sourceforge.net/projects/bbmap) and reads <50 bp post-trimming were discarded. Reads were aligned and quantified using STAR (2.6.0a) against the Ensembl human transcriptome (hg38.p12 genome release 96). Ensembl IDs were mapped to gene names and counts of genes with multiple IDs were aggregated. Low expression genes were removed if mean raw count <1.0 or mean counts per million (CPM) <1.0 for the dataset. Reads were normalized to CPM using the edgeR R package. Differential expression was calculated using the voom function in the limma R package. These data are available in the GEO database as GSE294453.

### Statistical analysis

Data are presented as mean ± standard error of the mean (SEM). Statistically significant differences, *p* values ≤0.05, were calculated using GraphPad Prism 6.0 statistical software and defined as * *p*≤0.05, ** *p*<0.01, *** *p*<0.001, and **** *p*<0.0001. *p* values were listed in each Fig. for non-significant trends.

### Data Availability Statement

Data, methods, and resources supporting the findings of this study which are not available in the manuscript or supplementary materials are available from the corresponding author, JKR, upon request. RNAseq gene expression data is publicly available through the GEO database. Mouse lung scRNAseq data have also been uploaded to the CELLxGENE data corpus as [*in progress*].

## RESULTS

### Chronic wound repair develops during metastatic outgrowth

#### Wound repair-related cells accumulate in the lung microenvironment during metastatic outgrowth

We hypothesized that alterations associated with large metastases, that are not present surrounding small metastases, develop during metastatic outgrowth. Therefore, to study how the metastatic microenvironment changes during outgrowth, we examined the lung directly adjacent to metastases of increasing size. By our definitions, small metastases in mice had a diameter of less than 150μm, which equates approximately to a cluster of 10 cells, and large metastases had a diameter of more than 300μm (Supplementary Fig. 1A). To put these sizes into perspective, only lung nodules larger than 5cm in diameter are clinically recommended for routine follow-up and the majority of these are found to be non-malignant [22]. Proportionately, human pulmonary nodules are approximately 150x larger than what we have defined as ‘large’ metastases in mice; however, the human lung, based on total lung capacity, is about 6000x larger than the mouse lung [23]. Thus, our ‘large’ metastases are particularly big when considering the actual size of mouse lungs compared to humans.

Our initial histological analysis of the lung metastatic microenvironment in MMTV-PyMT (mammary specific polyomavirus middle T antigen) transgenic mice showed an increase in cellularization in the adjacent lung during outgrowth (Fig. 2A), reminiscent of the epithelization that occurs during wound repair in the lung. We further observed an increase in the number of cells histologically identified as neutrophils and macrophages surrounding small versus large metastases (Supplementary Fig. 1C), cell types that are well-established contributors to wound repair responses. Consequently, to determine whether injury and repair occur in the lung adjacent to metastases during outgrowth, we examined whether cell types classically involved in lung wound repair are present in the lung surrounding metastases using cell type-specific markers. Normal wound repair in the lung is characterized by a cascade of distinct events involving neutrophils, macrophages, fibroblasts and AT2 cells. Using cell type specific immunohistochemical markers to stain serial sections from MMTV-PyMT metastatic lungs, we observed that the percentage of neutrophils (lymphocyte antigen 6G, Ly6G) surrounding metastases of increasing size remains consistent. The persistent presence of neutrophils throughout outgrowth is indicative of chronic wound repair, likely stalled during the inflammatory phase [24]. In contrast, there was a significant increase in the percentage of macrophages (EGF-like module-containing mucin-like hormone receptor-like 1, F4/80/Emr1), fibroblasts (fibroblast specific protein 1, Fsp1), and AT2 cells (prosurfactant protein C, proSP-C; Fig. 2A) surrounding large versus small metastases. Notably, these microenvironmental effects were highly localized and only found directly adjacent to metastases within 100μm of the metastatic perimeter. Very few differences were observed intratumorally (Supplementary Fig. 1D). Our analysis of wound repair-related cells in the lung surrounding metastases indicates that lung compaction, caused by growing metastases, is not solely responsible for what occurs in the lung during metastatic outgrowth suggesting that cells are actively proliferating and/or being recruited to the metastatic microenvironment (Supplementary Fig. 1E-G).

**Figure 2.**
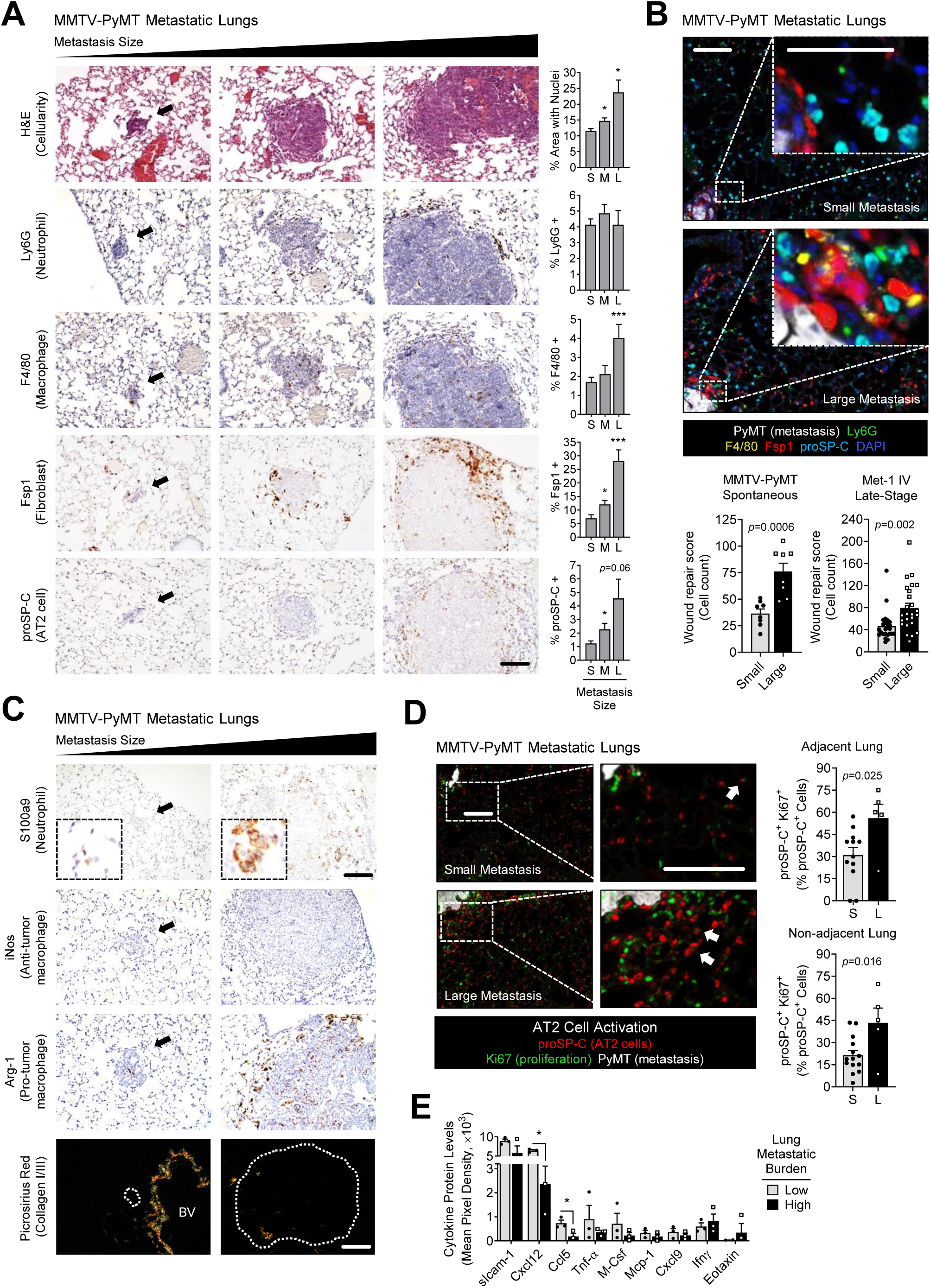
Chronic wound repair in the lung surrounding growing metastases. **A,** MMTV-PyMT metastatic lungs were H&E and IHC stained for cell-specific markers of lung wound repair (*n* >50 metastases from 2-7 mice per stain; see Supplementary Table 1 for sample number details). Shown are representative images from metastases of different sizes with black arrows indicating small metastases; scale bar = 100µm. The percentage of positively stained cells, normalized to the total number of cells, was quantified in the 100µm surrounding metastases. Mean ± SEM (unpaired *t*-tests with Welch’s correction when appropriate); * *p*≤0.05, *** *p*<0.001. S, small; M, medium; L, large metastases. **B,** Multispectral immunofluorescent staining for a lung wound repair cell signature in MMTV-PyMT and Met-1 metastatic lungs. Shown are representative images from MMTV-PyMT metastases of different sizes; scale bar = 25µm, inset zoom 3x. The wound repair score, calculated as the total number of wound repair-related cells in the 300µm surrounding metastases from the spontaneous MMTV-PyMT metastasis model (*n*=16 metastases from 2-4 mice; unpaired *t*-tests with Welch’s correction) and the late-stage Met-1 intravenous (IV) metastasis model (*n*=50 metastases from 5 mice; unpaired *t*-tests). Mean ± SEM. **C,** MMTV-PyMT metastatic lungs were stained for cell-specific activation markers. Shown are representative images from metastases of different sizes with black arrows indicating small metastases. For polarized light picrosirius red images, metastases are outlined in white. BV, blood vessel; scale bar = 100µm, inset zoom 4x. **D,** Multispectral immunofluorescent staining of MMTV-PyMT metastatic lungs for AT2 cell activation. Shown are representative images from metastases of different sizes with white arrows indicating proSP-C and Ki67 double-positive proliferating AT2 cells; scale bar = 25µm, inset zoom 3x. The percentage of proSP-C-positive cells that are Ki67-positive was quantified in the lung surrounding metastases: <300µm surrounding metastases for adjacent cells and >300µm for non-adjacent cells (*n*=17 metastases from 3-5 mice; unpaired *t*-tests). Mean ± SEM. S, small; L, large metastases. **E,** Cytokine array performed on lungs from mice with a low or high metastatic burden using the late-stage Met-1 IV metastasis model (*n*=3 mice per group; multiple unpaired *t*-tests with Welch’s correction). Mean ± SEM; * *p*≤0.05.

To determine the spatial distribution of wound repair-related cells in the evolving metastatic microenvironment, we developed a multispectral immunofluorescence (multi-IF) wound repair panel (Supplementary Fig. 1B). The data from this multi-IF panel reliably recapitulated data obtained from individual IHC stains (Fig. 2A and Supplementary Fig. 2A). By combining the counts for all cell types (neutrophils, macrophages, fibroblasts, and AT2 cells) a single ‘wound repair score’ was calculated to quantify wound repair for each metastasis. Higher wound repair scores were positively correlated with larger metastasis size (Supplementary Fig. 2B). In the MMTV-PyMT spontaneous metastasis model, wound repair is significantly increased in the lung adjacent to large metastases when compared to small metastases (Fig. 2B). Similar results were observed in a late-stage metastasis model in which MMTV-PyMT-derived Met-1 cells were intravenously (IV) injected into the tail veins of female FVB/NJ mice (Fig. 2B and Supplementary Fig. 2C). In this model mammary carcinoma cells directly seed the lung as opposed to spontaneous metastasis models in which mammary carcinoma cells disseminate from the primary tumor. Together these data indicate that wound repair-related cells consistently accumulate in the lung microenvironment during metastatic outgrowth, irrespective of the presence or absence of a primary tumor or pre-metastatic niche. Interestingly, a comparison of lung tissue surrounding metastases in spontaneous versus late-stage metastasis models indicated that while similar results were observed during metastatic outgrowth, there are significantly more F4/80-positive macrophages present in the lung surrounding small metastases in the late-stage model compared to the spontaneous metastasis model (Supplementary Fig. 2D). This suggests that lung pre-metastatic niche formation may play a role in the spontaneous metastasis model by priming alveolar macrophages toward a pro-tumor phenotype. Conversely, alveolar macrophages in the late-stage metastasis model, which have not been primed by a primary tumor, still attempt to clear particulates from the lung, including newly arrived metastatic cells. This multi-IF stain also validated the specificity of our cell type-specific markers. For instance, macrophages can express Fsp1/S100a4 depending on the context [25]. In our model there is little overlap between markers, including the macrophages (F4/80), fibroblast (Fsp1), and mammary tumor PyMT markers (Supplementary Fig. 2E). Finally, our data recapitulated prior studies showing a significant increase in neutrophils and fibroblasts during pre-metastatic niche preparation when comparing normal lungs to those with a low metastatic burden [26, 27], while also demonstrating that differences are present between the pre-metastatic (or low metastatic burden) and metastatic (high metastatic burden) niches (Supplementary Fig. 2F-G). These data suggest that changes to the microenvironment that arise during metastatic outgrowth are distinct from changes that develop during pre-metastatic niche formation and/or metastatic colonization.

#### Wound repair-related cells in the metastatic microenvironment become activated throughout metastatic outgrowth

Cell-specific functional markers were used to determine the activation status of wound repair-related cells within the metastatic microenvironment in MMTV-PyMT metastatic lungs. The calcium-binding protein S100a9 is a biomarker indicative of neutrophil activation/stimulation [28]. While total neutrophil numbers remain relatively constant during metastatic outgrowth, S100a9 levels are higher in neutrophils surrounding large versus small metastases (Fig. 2C). The increasingly punctate cellular localization of S100a9 in neutrophils throughout metastatic outgrowth is also indicative of increased neutrophils activation. While macrophage polarization and function within the tumor microenvironment is dynamic and extremely complex [29], the conventional markers used to identify anti-tumor and pro-tumor macrophage activity are inducible nitric oxide synthase (iNos) and arginase 1 (Arg-1), respectively. Macrophages within the lung metastatic microenvironment were almost exclusively iNos-negative and Arg-1-positive, and the number of Arg-1-positive cells is dramatically increased in the lung surrounding large versus small metastases (Fig. 2C and Supplementary Fig. 3A). The primary function of fibroblasts during wound repair is to secrete collagens. Using the picrosirius red stain for collagens I and III, we found that collagen deposition is low in the lung surrounding metastases, regardless of size (Fig. 2C and Supplementary Fig. 3B). We also examined levels of the traditional fibroblast activation marker alpha-smooth muscle actin (αSMA). Interestingly, very few αSMA-positive fibroblasts are present in the lung surrounding metastases (Supplementary Fig. 3C). The absence of collagen deposition and αSMA expression are additional indicators of chronic wound repair in the lung [30]. Finally, we examined AT2 cell activation, which is traditionally characterized by increased proliferation and trans-differentiation during wound repair resolution [31]. Lungs with metastases were multi-IF stained for the mammary tumor marker PyMT, the AT2-specific marker proSP-C, and the proliferation marker antigen kiel 67 (Ki67). AT2 cells surrounding large versus small metastases exhibited significantly more Ki67 staining, indicating increased AT2 proliferation/activation during metastatic outgrowth (Fig. 2D).

Next, we were interested to see whether large metastases influence the microenvironment surrounding other metastases within the same lung. Lungs with large metastases generally have a higher overall metastatic burden (including both large and small metastases) (Supplementary Fig. 3D), while low metastatic burden lungs exclusively have small metastases. Since AT2 proliferation was significantly higher in the non-adjacent lung as well as the adjacent lung microenvironments, we examined AT2 cell activation in the lungs surrounding small metastases from lungs with a low metastatic burden (S_L_) compared to small metastases in lungs with a high metastatic burden (S_H_). Interestingly, there were no significant differences in AT2 activation surrounding small metastases regardless of the overall metastatic burden (Supplementary Fig. 3E), suggesting that changes that develop in the microenvironment during outgrowth are highly specific to cells surrounding large metastases and not present throughout the entire lung.

#### A unique cytokine signaling signature is associated with metastatic outgrowth

Cytokines facilitate paracrine communication between cells [32]. To identify cytokines associated with metastatic outgrowth and metastasis-associated wound repair, we performed a cytokine array on lungs with low versus high metastatic burden. Whole lung lysates were collected one week following IV injection as a model for low metastatic burden lungs and three weeks post-injection as a model for high metastatic burden. Very few differences in cytokine levels were observed when comparing whole lungs with a low versus high metastatic burden, though there was an overall trend in lower cytokine levels with increasing metastatic burden (Supplementary Fig. 3F). There were two cytokines, CXC motif chemokine ligand 12 (Cxcl12) and CC motif chemokine ligand 5 (Ccl5), were significantly reduced in the lung during metastatic outgrowth (Fig. 2E).These cytokines are known to promote BC metastasis and recurrence [33, 34] and both play critical inflammatory roles in wound repair in the lung [35]. Interestingly, low levels of CXCL12 and CCL5 are associated with poor wound healing [36, 37]. Thus, our results represent an interesting overlap between paracrine signaling related to tumor cell growth and lung wound repair and suggest that cytokine signaling during metastatic outgrowth is unique, particularly when compared to what occurs during tumor cell dissemination and lung seeding. Importantly, within tissues, maximum cytokine diffusion is estimated to be around 250μm [38]. Since the whole lung was examined in this experiment, spatially restricted microenvironmental changes localized directly adjacent to metastases may have been difficult to detect. Nonetheless, these data suggest that chronic wound repair develops in the lung directly adjacent to metastases during outgrowth and that this process may be a unique metastasis-associated response.

### Metastatic outgrowth in the lung induces cell-specific changes in gene expression

To examine gene expression changes in the lung microenvironment in response to metastatic outgrowth, we performed single-cell RNA-sequencing (scRNAseq) on mouse lungs with low versus high metastatic burden using the Met-1 late-stage metastasis model. Magnetic resonance imaging (MRI) with cardiac- and breath-gating was performed to verify lung metastatic burden prior to tissue collection (Fig. 3A). Cellular transcriptomics data were compared to the published dissociation-related gene signature to confirm that the tissue dissociation protocol used was rapid and gentle enough to avoid significant gene expression changes [21]. We found that only 65 of 5504 cells had a dissociation signature greater than the established threshold of 7.5%, suggesting that our enzymatic dissociation technique had little effect on gene expression (Supplementary Fig. 4). We first examined organ-wide changes that develop during metastatic outgrowth in the lung by performing a bulk analysis, combining gene expression data from all cell types per condition, and found that peptide signaling, innate immunity, and inflammatory pathways were associated with high metastatic burden (Supplementary Fig. 5A and Supplementary Tables 2-3). To examine cell-specific gene expression changes associated with metastatic outgrowth, lung cells were divided into 26 clusters expressing unique gene expression signatures (Fig. 3B). Metastatic mammary tumor cells were identified using the *PyMT^+^CyclinD1^+^* gene signature [39], while general cell types were determined by examining the top 3-5 genes per cluster (Supplementary Fig. 5B-C and Tables 1-2). Gene expression analysis identified 8 clusters of common cell types (Fig. 3C and Tables 3-4). Significant gene expression changes, associated with high versus low metastatic burden were observed in dendritic, endothelial, lymphocyte, stromal, monocyte/ macrophage, and epithelial cell populations (Fig. 3D). These data illustrate how dramatically the metastatic microenvironment is altered during metastatic outgrowth.

**Figure 3.**
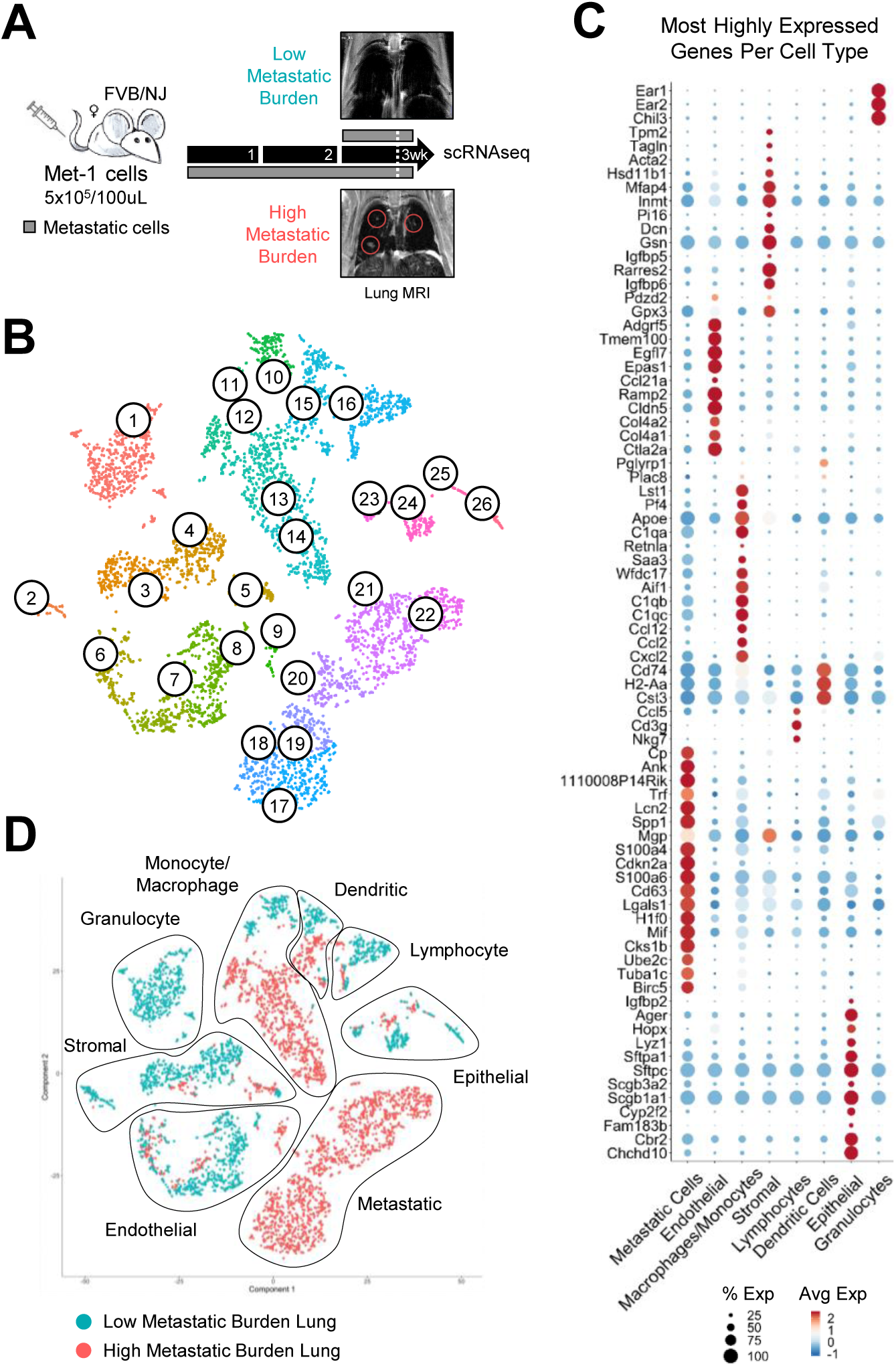
Cell type-specific gene expression within the lung during metastatic outgrowth. **A,** Schematic of single cell RNA-sequencing (scRNAseq) experimental design using the late-stage Met-1 metastasis model. To verify metastatic burden, lungs from mice with a low or high metastatic burden (n=1 mouse per group) were scanned by magnetic resonance imaging (MRI) 3 days prior to tissue collection. Shown are representative coronal MRI images of lungs with low or high metastatic burden, with large metastases circled in white. Lungs were then collected from perfused mice and enzymatically dissociated for downstream scRNAseq. **B,** t-SNE visualization of lung cells clustered by gene expression. **C,** Individual cell clusters were grouped by cell type depending on the top 3-5 most highly expressed genes per cluster, using known cell type-specific genetic markers. Average expression (Avg Exp) was defined as average log fold change between one cell type group and all other cell types. Percent expression (% Exp) was defined as the percentage of cells within a cell type group that express each gene. **D,** t-SNE visualization of cell clusters grouped by cell type and colored by metastatic burden.

### Metastatic outgrowth promotes differential gene expression in lung AT2 cells

Lung epithelial cells are the most numerous cells within the lung and are likely to interact extensively with metastatic cells [13]. We identified distinct cell types within the epithelial cell cluster using the top 3-5 genes per cluster (Fig. 4A). The AT2 cell population, cluster 24, was verified by examining surfactant protein gene expression (Supplementary Fig. 6A-B). Of the 4 lung epithelial cell clusters, AT2 cells showed the most pronounced shift in gene expression in lungs with a high versus low metastatic burden (Fig. 4B). During metastatic outgrowth, AT2 cells upregulated the levels of genes encoding factors involved in wound repair and proliferation and downregulated genes involved in apoptosis and surfactant production (Fig. 4C, Table 5, and Supplementary Table 4). This metastasis-associated gene signature is suggestive of an activated AT2 phenotype in which the cells have shifted their functional focus from surfactant production towards a wound repair-related phenotype. Many of these genes, in other cell types, are known to contribute to breast cancer and metastatic progression (Supplementary Table 4).

**Figure 4.**
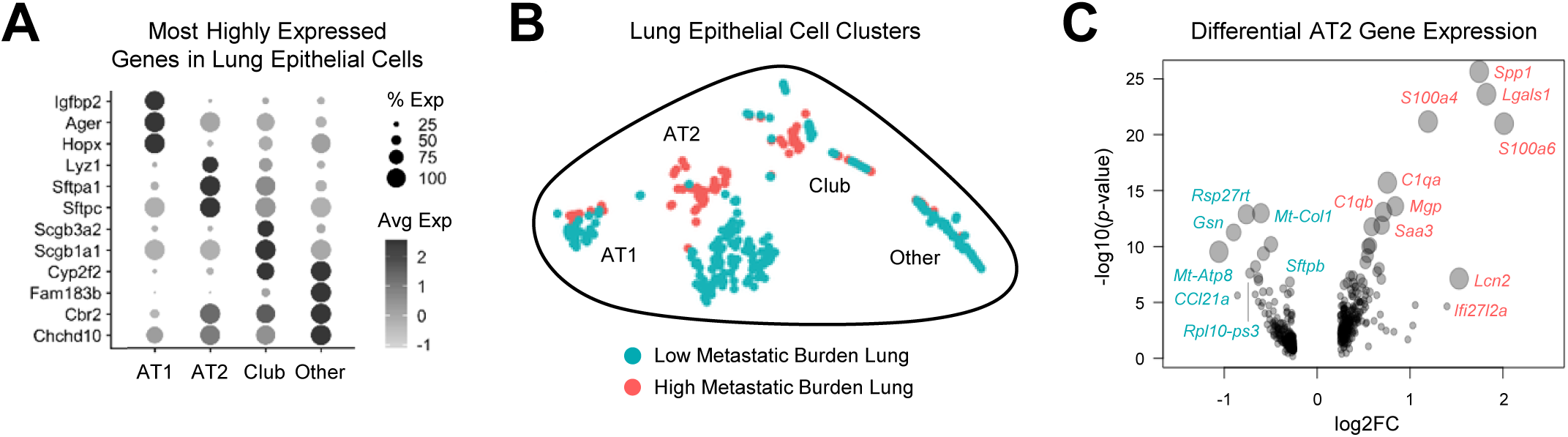
Lung AT2 cell gene expression during metastatic outgrowth. **A,** Epithelial cell clusters grouped by cell type depending on the top 3-5 most highly expressed genes per cluster, using known cell type-specific genetic markers. Average expression (Avg Exp) was defined as average log fold change between one cell type group and all other cell types. Percent expression (% Exp) was defined as the percentage of cells expressing each gene. Club cells, also known as Clara cells, are bronchiolar epithelial cells. **B,** t-SNE visualization of lung epithelial cells clustered by gene expression and colored by metastatic burden. **C,** Differential gene expression in AT2 cells from lungs with a low versus high metastatic burden. Gene expression displayed by fold change (FC) and *p*-value. Circle size represents the significance of differential gene expression. Genes labelled in pink were enriched in AT2 cells from lungs with a high metastatic burden, while genes labelled in blue were enriched in AT2 cells from lungs with a low metastatic burden.

### Paracrine interactions with BC cells stimulate AT2 activation

To directly investigate how BC cells affect AT2 cell activation and function (Fig. 5A), we used three TNBC cells lines (SUM159PT, BT549, and MDA453) and two AT2 cell models (aged A549 cells and differentiated induced pluripotent stem cells (iPSC), iAT2 cells). A549 cells are an immortalized lung carcinoma cell line originating from AT2 cells. When cultured long-term (sub-cultured post-thaw more than 8 times), A549 cells regain a more AT2 cell-like phenotype with slowed growth and increased expression of AT2-specific surfactant proteins (*SFTP*) (Supplementary Fig. 6C) [17]. iAT2 cells are created by differentiating iPSC cells through the endoderm stage to AT2 cells (Supplementary Fig. 6D-E) [17]. Since our in vivo data suggest that BC and AT2 cells interact primarily through localized paracrine mechanisms, we used a no-contact co-culture method to study the indirect interactions between these cell types (Fig. 5B). Co-culture with BC cells significantly increased AT2 cell growth compared to AT2 cells cultured alone, as measured by a cell viability assay (Fig. 5C). H&E staining of AT2 cells co-cultured with BC cells, when compared to AT2 cells cultured alone, exhibited increased size and an accumulation of intracellular vacuoles (Fig. 5D). Similar changes in cell morphology were observed in AT2 cells treated with conditioned media (CM) from SUM159PT cells (Supplementary Fig. 6F). These vacuoles strongly resembled AT2-specific lamellar bodies. To investigate the identity of these vacuoles, iAT2 cells, cultured alone or co-cultured with BC cells, were stained with lysotracker to examine lamellar body levels. The distribution of lamellar body stain was diffuse in iAT2 cells cultured alone, while co-culture with TNBC cells caused a more punctate distribution of staining (Fig. 5E). Quantification of this stain showed that co-culture with BC cells promotes lamellar body numbers and/or size in iAT2 cells (Fig. 5F). Whether this represents differential surfactant production or changes to surfactant secretion in AT2 cells remains unclear. It does, however, indicate a shift in AT2 cell function from its primary role facilitating breathing toward a more wound repair-related phenotype. To better understand how AT2 cells are affected by BC cell-derived secreted factors, we performed bulk RNAseq on iAT2 cells cultured alone or co-cultured with SUM159PT or MDA453 cells (Supplementary Fig. 6G and Table 6). Gene expression in human iAT2 cells (cultured alone, 509 genes) was compared to scRNAseq data from mouse AT2 cells (low metastatic burden lungs, 194 genes). Little overlap was observed between these two datasets (25 genes; Supplementary Fig. 6H and Supplementary Table 5). Similarly, very little overlap was observed in differentially expressed genes in human iAT2 cells (co-cultured with BC cells vs cultured alone) and those in mouse AT2 cells (high metastatic burden vs. low metastatic burden lungs) (5 genes; Supplementary Fig. 6I and Supplementary Table 5). Functionally, however, several biological pathways overlapped between these data sets, and one pathway in particular, secreted/signaling peptides, was enriched in all three AT2-BC interaction datasets (Fig. 5G, Table 7, and Supplementary Tables 6-7). These data indicate that paracrine interactions between BC and AT2 cells induce an activated AT2 phenotype characterized by amplified cell growth, altered cell morphology, increased lamellar body numbers, and differential expression of secreted factors (Table 8). Functionally, these data suggest that during metastatic outgrowth AT2 cells are capable of dramatically influencing the behavior of surrounding cell populations within the lung.

**Figure 5.**
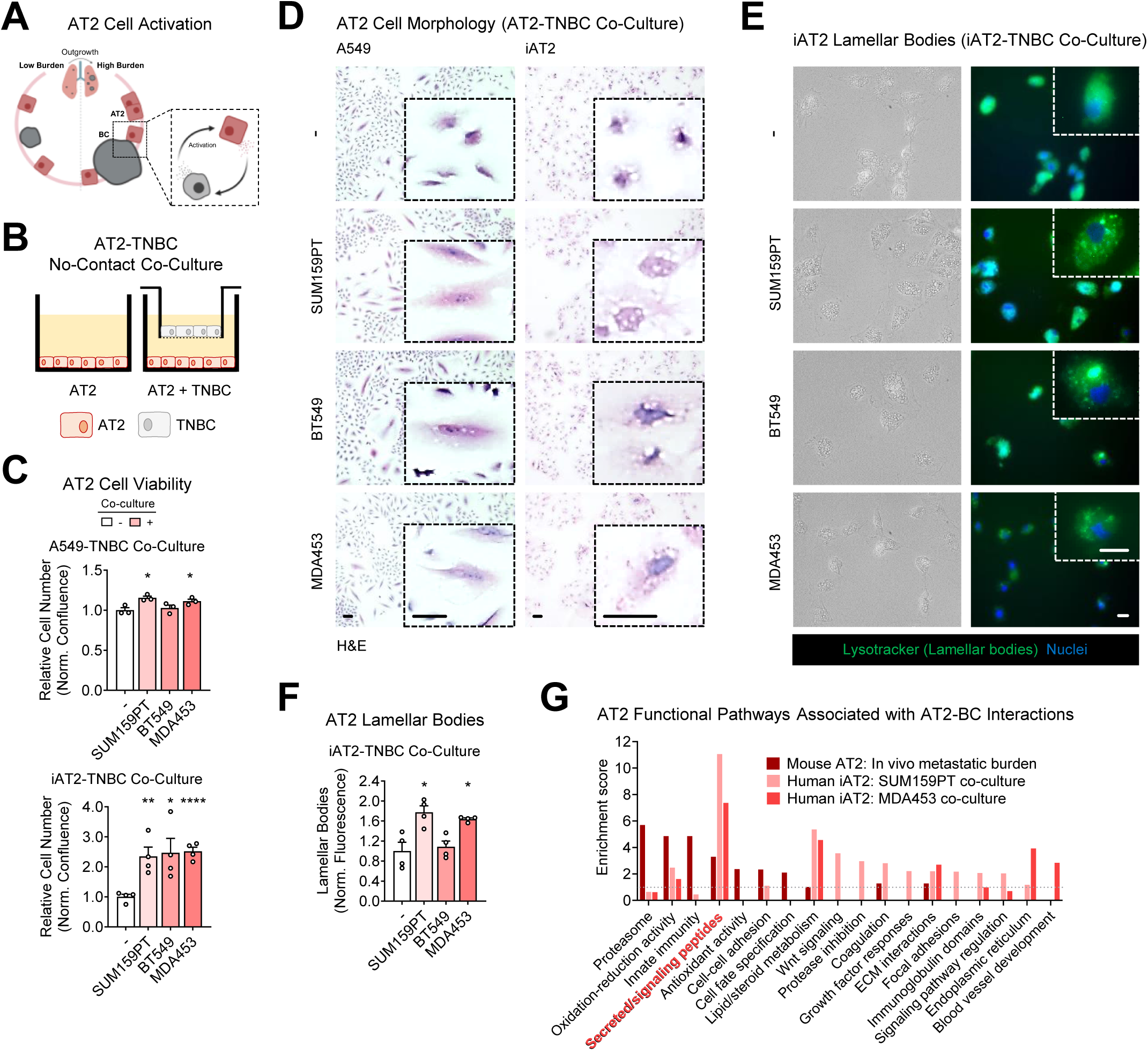
AT2 cell activation by BC cells. **A,** Model indicating the cellular interactions being interrogated: BC activation of AT2 cells. **B,** Schematic of no contact co-culture experimental design used to investigate the effects of TNBC cells on lung AT2 cells. **C,** AT2 cell viability was measured by crystal violet assay following co-culture with TNBC cells: 5 day co-culture for A549 cells and 7 day for iAT2 cells. Relative cell numbers were normalized to the mean confluence of AT2 cells cultured alone (-). Mean ± SEM (unpaired *t*-tests with Welch’s correction when appropriate); * *p*≤0.05, ** *p*<0.01, **** *p*<0.0001. **D,** AT2 cell morphology was examined by H&E staining following co-culture with TNBC cells for 3 days. Control cells were cultured alone (-). Shown are representative images of cell size and shape; scale bar = 10µm, inset zoom 4x for A549 cells and 6x for iAT2 cells. **E,** iAT2 lamellar bodies were imaged using lysotracker staining following co-culture with TNBC cells for 3 days. Control cells were cultured alone (-). Shown are representative images of lamellar body size and shape; scale bar = 10µm, inset zoom 3x. **F,** iAT2 lamellar body levels were measured by quantifying lysotracker staining in iAT2 cells co-cultured with TNBC cells for 3 days. Lamellar body fluorescence was normalized to cell number and the mean fluorescence of iAT2 cells cultured alone (-). Mean ± SEM (unpaired *t*-tests with Welch’s correction when appropriate); * *p*≤0.05. **G,** Bulk RNAseq was performed on RNA collected from iAT2 cells co-cultured with SUM159PT and MDA453 cells for 5 days, while control iAT2 cells were cultured alone. Function biological pathways in AT2 cells were determined using differentially expressed gene lists from mouse AT2 cells in metastatic lungs and human iAT2 cells co-culture with TNBC cells. The enriched pathways from these datasets were then compared to identify overlapping biological pathways.

### AT2 secreted factors promote BC cell growth

Modified secreted factor gene expression in AT2 cells alludes to the possibility that these cells reciprocally affect BC cell function (Fig. 6A). A complementary no-contact co-culture experiment was performed to investigate the effects of AT2-derived secreted factors on BC cell viability (Fig. 6B). Co-culture with AT2 cells significantly increased TNBC cell growth compared to TNBC cells culture alone, as measured by a cell viability assay (Fig. 6C). While the TNBC cell viability fold changes were relatively small, since TNBC cell lines have extremely high baseline proliferative rates with doubling times of between 1.0 - 2.5 days, depending on the cell line tested [40], any increase in proliferation is biologically noteworthy. These data suggest that AT2 cells promote BC proliferation within the lung.

**Figure 6.**
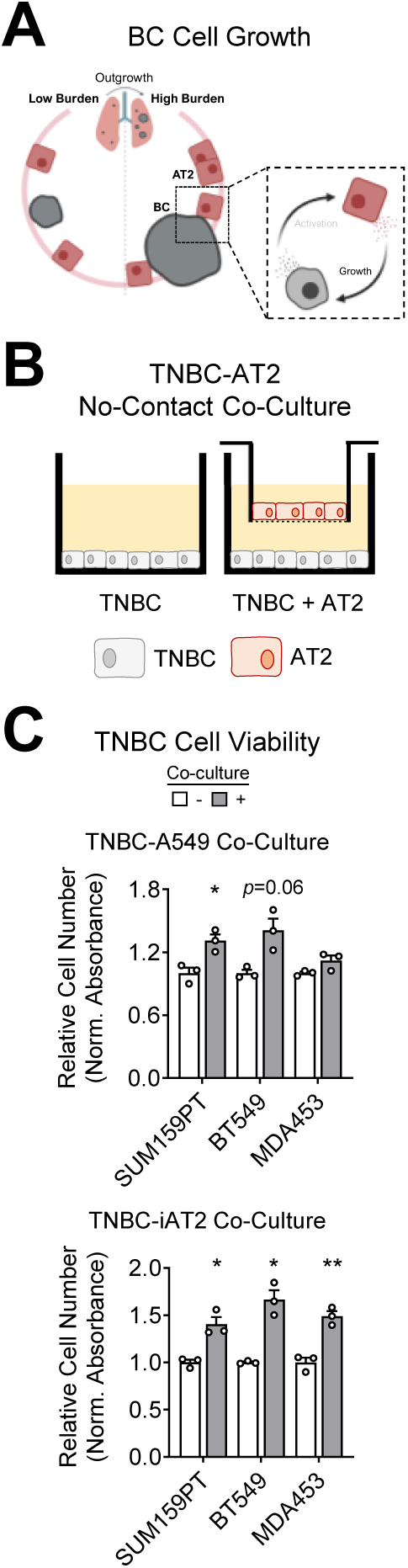
BC cell viability in response to AT2 secreted factors. **A,** Model indicating the cellular interactions being interrogated: AT2 induction of BC cell growth. **B,** Schematic of no-contact co-culture experimental design used to investigate the effects of lung AT2 cells on TNBC viability. **C,** TNBC cell viability was measured by crystal violet assay following co-culture with AT2 cells: 5 day co-culture for A549 cells and 7-8 day for iAT2 cells. Relative cell numbers were normalized to the mean absorbance of TNBC cells cultured alone (-). Mean ± SEM (multiple unpaired *t*-tests with Welch’s correction); * *p*≤0.05, ** *p*<0.01.

### PDE4 inhibition prevents AT2-BC reciprocal activation

Overall, our data indicate that throughout metastatic outgrowth, and the development of metastasis-associated wound repair, AT2 cells become activated in the presence of BC cells and, subsequently, AT2 cells reciprocally promote BC cell growth through indirect paracrine signaling. We hypothesized that interrupting the reciprocal paracrine interaction between AT2 and BC cells may inhibit lung metastatic outgrowth (Fig. 7A). To identify a suitable target for pharmaceutical intervention we examined predicted upstream regulators of differentially expressed secreted factors identified in AT2 cells from high metastatic burden mouse lungs and TNBC co-cultures. We focused primarily on transcriptional regulators with established treatment strategies that could easily be repurposed for use in patients with BC. Notably, a large percentage of BC-induced AT2 secreted factors, identified in RNAseq datasets from both human and mouse cells, are known cAMP response element-binding protein (CREB)-regulated genes (Fig. 7B, Supplementary Fig. 7A, and Table 8) [41]. CREB was further identified as a potential upstream regulator of AT2 secreted factor genes in all three datasets by Ingenuity Pathway Analysis (mouse metastatic lungs *p*=9.77E-05, SUM159PT co-culture *p*=3.93E-03, and MDA453 co-culture *p*=3.12E-03) [42].

**Figure 7.**
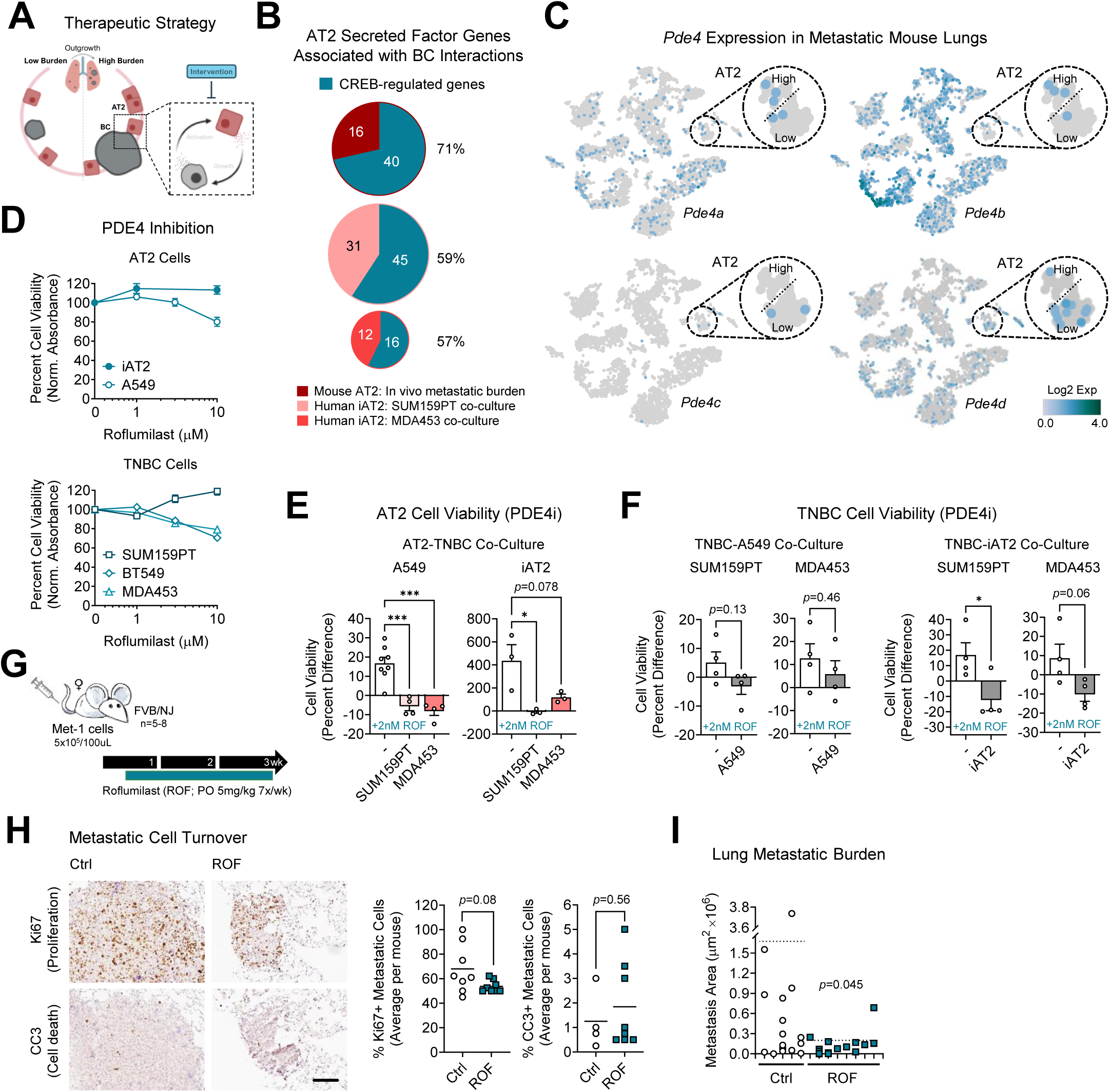
Targeting CREB-regulated AT2 secreted factors to block AT2-BC interactions and metastatic outgrowth. **A,** Model indicating the cellular interactions being interrogated: targeting AT2-BC interactions during metastatic outgrowth. **B,** The proportion of AT2 secreted factor genes enriched following BC interactions that are CREB-regulated for all three AT2 RNAseq datasets, percentage indicated. **C,** t-SNE visualizations of cells from metastatic mouse lungs clustered by gene expression and colored by *Pde4* (phosphodiesterase 4) isoform gene expression. **D,** Percent cell viability was measured by crystal violet assay after a 3 day treatment with vehicle DMSO (0) or increasing concentrations of the PDE4 inhibitor roflumilast. Data was normalized to the mean absorbance of DMSO-treated cells; mean ± SEM. **E,** AT2 cell viability was measured by crystal violet assay following co-culture with TNBC cells and treatment with 2nM roflumilast (ROF): 5 days for A549 cells and 7 days for iAT2 cells. The percent difference in cell viability, between DMSO and 2nM ROF treated cells, was calculated for AT2 cells cultured alone (-) or co-culture with TNBC cells. Mean ± SEM (one-way ANOVA with Tukey’s multiple comparison test); * *p*≤0.05, *** *p*<0.001. **F,** TNBC cell viability was measured by crystal violet assay following co-culture with AT2 cells and treatment with 2nM ROF: 5 days for A549 cells and 7 days for iAT2 cells. The percent difference in cell viability, between DMSO and 2nM ROF treated cells, was calculated for TNBC cells cultured alone (-) or co-culture with AT2 cells. Mean ± SEM (unpaired *t*-tests); * *p*≤0.05. **G,** Schematic of metastatic outgrowth experimental design using the late-stage Met-1 metastasis model (*n*=5-8 mice per group). Met-1 cells were IV injected into female mice and oral treatment with 5mg/kg ROF began 3 days later. Mice were treated daily for 3 weeks and metastatic lung tissue was collected. **H,** Lungs were stained for cell turnover markers Ki67 and cleaved-caspase 3 (CC3) by IHC. Shown are representative images of lung metastases; scale bar = 100µm. The percentage of positively stained cells was scored per metastasis and averaged per mouse; mean (unpaired *t*-tests with Welch’s correction when appropriate). **I,** Metastatic lungs were stained for the Met-1 mammary-specific marker PyMT. Metastatic burden was quantified in serial sections as the area of PyMT-positive metastases, means per group are indicated as dotted lines (unpaired *t*-test). Representative data is shown from a single section indicating the number and size of metastases per mouse; each notch on the x-axis represents an individual mouse.

CREB is the transcription factor responsible for regulation of genes associated with the cyclic adenosine monophosphate (cAMP) – protein kinase A (PKA) – CREB signaling axis. cAMP-PKA-CREB signaling regulates a number of physiological processes and aberrant signaling plays a role in a variety of cancers [43]. cAMP has been specifically linked to cytokine secretion in lung epithelial cells [44], and CREB is considered to be a key regulator of BC metastasis [45]. cAMP-PKA-CREB signaling, which has an overall anti-inflammatory effect, is negatively regulated by phosphodiesterase (PDE) activity through the hydrolysis of cAMP [46]. PDE4 is one of several PDEs in the lung, where it is ubiquitously expressed and collectively contributes to organ-wide inflammatory responses [47]. Notably, PDE4 is the primary PDE expressed in AT2 cells [48]. In normal, healthy lung tissue and AT2 cells from humans and mice, *PDE4* isoforms (*PDE4A-D*) are expressed at relatively low levels (Supplementary Fig. 7B) [49]. In mouse lungs with metastases, *Pde4* isoforms are expressed at various levels in many cells types (Fig. 7C). During metastatic outgrowth, isoform expression in AT2 cells shifts from *Pde4c* and *Pde4d* in lungs with a low metastatic burden toward *Pde4a* and *Pde4b* in lungs with a high metastatic burden, though these changes are observable trends and not statistically significant (Fig. 7C). Overall, these data suggest that PDE4 inhibition may be a rationale method for targeting reciprocal interactions between AT2 and BC cells.

Non-steroidal anti-inflammatory PDE4 inhibitors are commonly used to treat pathological lung conditions. Roflumilast (ROF) is a routinely used second-generation oral PDE4 inhibitor FDA-approved for the treatment of chronic obstructive pulmonary disease (COPD). Dose-response cell viability experiments were performed to test the sensitivity of AT2 and TNBC cells to increasing doses of ROF. While some statistical differences were observed with varying doses of ROF, none of the effects illustrated biologically relevant effects on cell viability in AT2 or TNBC cells (cell viability following 10µM ROF ranging from 71-119%; Fig. 7D), especially considering physiologically relevant doses of ROF are between 1-2nM [50]. Similar results were observed in estrogen receptor-positive BC cells treated with ROF (Supplementary Fig. 7C). Likewise, the sensitivity of AT2 and TNBC cell viability was examined in response to increasing doses of another second-generation PDE4 inhibitor cilomilast (CILO) and no biologically relevant effects on cell viability were observed (Supplementary Fig. 7D). Met-1 mouse mammary carcinoma cells also showed little response to PDE4 inhibition, consistent with what we observed in human BC cells (Supplementary Fig. 7E). These data indicate that AT2 and BC cells are resistant to ROF when cultured alone.

To test whether BC-induced activation sensitizes AT2 cells to PDE4 inhibition, cell viability was examined following simultaneous treatment with conditioned media (CM) from TNBC cells and ROF. A549 cells were sensitized to ROF when activated by secreted factors from TNBC cells, as measured by reduced cell viability compared to A549 cells cultured in normal media (Supplementary Fig. 7F). Similarly, since our data indicate that AT2 and TNBC cells reciprocally promote growth when cultured in no-contact co-culture conditions (Fig.s 5C and Fig. 6C), we tested whether reciprocal activation may be necessary to sensitize AT2 and TNBC cells to ROF. To investigate this, AT2 and TNBC cell viability was quantified following 2nM ROF treatment with or without co-culture. When cultured alone, 2nM ROF marginally promoted cell growth in several cell lines (Supplementary Fig. 7G). Interestingly, AT2-TNBC paracrine interactions reversed these ROF effects. ROF inhibited cell viability in AT2 cells co-cultured with TNBC cells (Fig. 7E). Similarly, TNBC cell viability was inhibited by ROF only when co-cultured with AT2 cells (Fig. 7F). These data suggest that reciprocally activated AT2-TNBC cells respond to PDE4 inhibition.

### PDE4 blockade limits lung metastatic outgrowth

PDE4 inhibition, using ROF, as a treatment for BC patients with lung metastases is a promising therapeutic strategy. Not only does our data suggest that ROF impedes the pro-growth effects of AT2 cells on BC cells during metastasis, it also indicates that chronic wound repair develops during metastatic outgrowth which is reminiscent of the chronic inflammation present in patients with COPD who are routinely prescribed ROF. Thus, we tested the effectiveness of ROF in blocking metastatic outgrowth using a late-stage metastasis mouse model of BC. Once Met-1 cells colonized the lungs, 3 days following tail vein injection, mice were administered oral ROF daily at a dose of 5mg/kg for 3 weeks (Fig. 7G). Daily treatment with ROF was well tolerated with no changes in mouse weight over the course of the experiment (Supplementary Fig. 7H). To examine the tumor cell response to ROF, we stained metastatic lungs by IHC for proliferation and cell death markers, Ki67 and cleaved-caspase 3 (CC3) respectively. While the data did not reach statistical significance, there is a clear trend toward reduced metastatic proliferation in ROF treated lungs with little to no change in cell death (Fig. 7H). Metastatic burden was analyzed by IHC staining lung serial sections for the Met-1 mammary-specific marker PyMT. While ROF treatment had no effect on the number of metastases, it did significantly reduce the size of lung metastases (Fig. 7I and Supplementary Fig. 7I) suggesting that ROF inhibited outgrowth of established lung metastases rather than killing metastatic cells. Thus, our pre-clinical analysis suggests that PDE4 may be a viable target in BC patients with lung metastasis.

## DISCUSSION

When defining the stages of the metastatic cascade many studies and reviews end the cascade at the stage of colonization. Outgrowth is a phase that is often overlooked as a distinct stage of metastasis and understudied in preclinical models. However, outgrowth of metastatic disease significantly impacts patient survival and quality of life. Our study shows that widespread cell-specific changes occur during metastatic outgrowth within the lung (Fig. 3). During outgrowth, unique, critical interactions between cancer cells and resident cells within the metastatic tissue occur that support progression to overt metastasis. This study identified targetable reciprocal interactions that may lead to treatment strategies for patients with BC lung metastases.

Growing metastases cause damage to surrounding tissue within the lung leading to a chronic wound repair response. Interestingly, our data indicates that while there are many similarities to classic wound repair, metastasis-associated wound repair is a unique process likely caused by a combination of physical injury and cancer cell paracrine signaling (Fig. 1-2). AT2 cell function is significantly altered during metastatic outgrowth (Fig. 3-4). Our data is consistent with previous studies noting the potential importance of AT2 cells in pre-metastatic niche formation and metastatic colonization. We expanded on those studies by examining how AT2 cells change throughout BC metastatic outgrowth and how they reciprocally support metastatic progression (Fig. 5-6).

Several functional biological pathways were altered in AT2 cells following interaction with BC cells, and our investigation focused on the commonly enriched secreted/signaling peptides pathway as a potential mechanism for how AT2 cells promote TNBC cell viability. We identified several CREB-regulated AT2 secreted factors involved in inflammation, immune responses and chemotaxis which could dramatically impact cell behavior within the surrounding microenvironment (Supplementary Table 7). PDE4 inhibition in lung epithelial cells reduces inflammatory/immune secreted factor levels and has previously been shown to enhance resolution of inflammation in the lung [51]. By inhibiting PDE4 activity, we successfully reduced reciprocal AT2-BC cell activation and metastatic outgrowth (Fig. 7). Importantly, AT2 cells play a critical role in regulating immune response within the alveolus through integration of multi-cellular signals. Thus, changes to AT2 gene expression and function could cause a cascade of effects within the lung microenvironment with a collective anti-metastatic effect. This effect might also be complemented by the anti-inflammatory consequences of PDE4 inhibition on other cells within the lung, where PDE4 is ubiquitously expressed [47].

Several PDE4 inhibitors are FDA-approved to treat inflammatory conditions, but ROF is the only PDE4 inhibitor currently approved for the treatment of COPD. For over a decade, COPD patients have been prescribed a daily oral dose of 500µg ROF to counter the long-term inflammatory consequences of COPD [50]. ROF is generally well-tolerated [52], and the safety of extended daily treatment is firmly established [53]. There is some evidence suggesting that ROF may be associated with an increased incidence of certain cancers [54], but other studies have shown its effectiveness as an anti-cancer therapy, particularly when given in combination with chemotherapy [55, 56]. Our data strongly suggest that ROF may serve as a viable lung metastasis-specific treatment strategy, but future studies will investigate its overall anti-tumor effects in BC, particularly when combined with standard-of-care chemotherapy. Future studies will also investigate its effects on the early stages of metastasis, given the importance of AT2 cells in pre-metastatic niche preparation and metastatic colonization, as well as hormone-related BC. Finally, since ROF caused little-to-no cell death of metastases, future studies need to determine whether the growth inhibitory effects of ROF are reversible once drug treatment is discontinued. Determining whether ROF induces irreversible cellular senescence or reversible quiescence/dormancy will help to pinpoint which anti-cancer drugs will be most effective when combined with ROF.

PDE4 inhibitor development is proceeding at a rapid pace. ROF is a non-specific PDE4 inhibitor that blocks activity of all PDE4 subtypes with a high affinity for PDE4B and PDE4D. Our interrogation of publicly available scRNAseq data indicated that baseline levels of *PDE4* isoforms are different between human and mouse lungs and AT2 cells. Mouse lungs predominantly express *Pde4b*, while human lungs and AT2 cells express relatively high levels of *PDE4D* (Supplementary Fig. 7B). Isoform specific inhibitors are under development and may prove to be more effective against lung metastases. Additionally, inhaled PDE4 inhibitor therapy is being investigated to directly target inflammation in the lung [57]. The FDA recently approved nebulizer delivered ensifentrine, a dual PDE3/PDE4 inhibitor, for the treatment of adults with COPD [58]. Since many of the effects observed in our studies are localized to the lung directly adjacent to metastases, inhaled administration of drugs may be a better way to target focal microenvironmental changes induced by growing metastases. Future studies are needed to clarify the most efficacious mode of PDE4 targeted therapeutic intervention for patients with lung metastases. Ultimately, this study indicates that targeting metastasis-associated wound repair and AT2 activation during lung metastatic outgrowth may prove to be a promising avenue for treatment of metastatic BC.

## ACKNOWLEDGMENTS

Financial support was provided by several grants, including: R01 CA258317 (JKR), NIH T32CA190216-01A1 (JLC), ACS IRG 16-184-56 (JLC), METAvivor Early Career Investigator Award 25B0307 (JLC).The authors appreciate the contribution to this research made by E. Erin Smith, HTL (ASCP)CM QIHC; Allison Quador, HTL(ASCP)CM; and Jessica Arnold HTL (ASCP)CM of the University of Colorado Cancer Center Pathology Shared Resource. This resource is supported in part by the Cancer Center Support Grant (P30CA046934). This study was also partly supported by the National Institutes of Health (P30CA046934) by utilizing the Bioinformatics and Biostatistics Shared Resource. Bioinformatics analyses were supported by the Alpine HPC system, which is jointly funded by the University of Colorado Boulder, the University of Colorado Anschutz, Colorado State University, and the National Science Foundation (award 2201538). For MRI-based animal imaging the authors would like to acknowledge the NIH SIFAR shared instrumentation grant S10OD027023. Finally, the authors would like to thank bioinformatician G. Devon Trahan for his assistance with scRNAseq analysis and honor the memory of veterinary pathologist Linda Kassenbaum Johnson, DVM, for assistance with histological scoring of mouse lung tissue.

