## Supplementary Methods for "Metastasis-associated wound repair promotes reciprocal activation of the lung epithelium and breast cancer metastases during outgrowth"

### SUPPLEMENTARY MATERIALS AND METHODS

**Cell culture.** MCF7 estrogen receptor-positive (ER+) BC cells were purchased from ATCC in 2012 and maintained in MEM with 5% FBS, 1% non-essential amino acids (NEAA), and 6ng/mL insulin. T47D ER+ BC cells were kindly provided by Kate Horwitz (University of Colorado) in 2009 and maintained in DMEM with 10% FBS. 66Cl4 mouse mammary carcinoma cells [1, 2] were kindly provided in 2017 by Heide Ford (University of Colorado) with permission granted by Fred R. Miller (Wayne State University). 66Cl4 cells were maintained in DMEM with 10% fetal calf serum (FCS), 1% L-glutamine, and 1% NEAA.

Conditioned media (CM) treatments. Conditioned media was collected from TNBC cells cultured in 10cm plates for 3 days. Control media was cultured in empty plates for 3 days. Media was collected and cells per plate counted. Media was centrifuged at 1,500 rpm for 4min to remove cellular debris, filter sterilized and stored at -80ºC. To test the effects of TNBC secreted factors on AT2 cells, a 1:1 mixture of AT2 media to TNBC CM was added to AT2 cells.

**In vivo mouse experiments.**

Lung MRI scans. Each mpMRI session consisted of a tri-pilot localizer, followed by gated fast spin echo proton density weighted PD-RARE (Rapid Acquisition with Refocusing Echoes) sequences in axial and coronal planes using a 38-mm mouse body RF coil. The following sequence parameters were used: field of view 30x30 mm; slice thickness 1 mm; number of slices 20 (coronal) and 24 (axial); repetition time TR=2200 ms; echo time TE=32 ms; RARE factor 8; number of averages 4; matrix size 320x320. Total acquisition time for all sequences was 11 min 6 sec. MRI image acquisition and analysis was performed using Bruker ParaVision 360neo v2.0 software by a MRI physicist with >15 years of experience.

**Histology.**

IHC Metastasis size. Size was determined using ImageJ software. Each metastasis was circled to compute area in pixels$^2$. Metastasis diameter in μm was calculated using the conversion factor 1.25 ImageJ pixels = 1 μm and the following calculations:

$$\text{metastasis area } (\mu m^2) = \left(\frac{\sqrt{\text{metastasis area in pixels}^2}}{1.25}\right)^2$$

$$\text{metastasis diameter } (\mu m) = \left(\sqrt{\frac{\text{metastasis area}}{\pi}}\right) \times 2$$

When comparing small, medium and large metastases: small metastases had a diameter (d) <150μm, medium metastases d=150-300μm, and large metastases d>300μm. When comparing small versus large metastases: small metastases had d<200μm and large metastases d>200μm. ImageJ was also used to quantify IHC staining in the adjacent 100μm of lung surrounding metastases (Supplementary Figure 1a). Data was presented as the percentage of positively stained cells relative to the total number of cells.

Lung compaction investigation. This analysis compared cellular content in the same total area of the lung surrounding metastases regardless of metastasis size (excluding metastatic and intratumoral cells). The 100μm surrounding the largest lung metastasis in the MMTV-PyMT lung cohort, with a radius of 325μm, was used to determine the total area being compared surrounding all metastases, ~$5.7 \times 10^5$ $\mu m^2$ (radius of 425μm from the center of each metastasis). ImageJ was used to quantify cell counts in the lung surrounding metastases. The cellular count from our smallest metastases, with a radius of 25μm, was used as a reference for comparison. Data was presented as the number of cells normalized to the cell count for the smallest metastasis (Supplementary Figure 1e).

Metastatic cell turnover. Ki67 and cleaved-caspase 3 (CC3) IHC levels in mouse lung metastases were scored visually by an experienced histotechnician. The percentage of positively stained cells was determined for each metastasis. Data is presented as the average staining in metastases per mouse.

Multispectral immunofluorescence (multi-IF). Only ~1-2% of each image included metastatic cells; the remaining image area was focused on surrounding lung tissue and vasculature, ~96% and ~4% respectively (Supplementary Figure 1b). Metastasis size was determined using Phenochart software (Akoya Biosciences). The width and height (μm) of each metastasis was measured and averaged to calculate metastasis diameter. When comparing small versus large metastases: small metastases had $d<200\mu m$ and large metastases $d>200\mu m$.

Metastatic microenvironment analysis. Basic mathematical equations for a circle (listed below) were used in multi-IF analyses to determine: 1) the area adjacent to metastases, within a radius of ≤300μm, and 2) whether individual cells were located adjacent or non-adjacent to metastases (Supplementary Figure 1b). A comprehensive wound repair score was calculated by combining the counts for all cell types (neutrophils, macrophages, fibroblasts, and AT2 cells) within the lung adjacent to metastases. Data are presented as the number of positive cells normalized to area.

$$A = \pi r^2$$

$$d = r \times 2$$

$A$ = area

$r$ = radius

$d$ = diameter

$$r^2 = (x - h)^2 + (y - k)^2$$

$(h, k)$ = location of the center of the circle (corner of the ROI with the metastasis)

$(x, y)$ = location of the cell of interest

**Bioinformatics**

Enzymatic dissociation of lungs for scRNAseq. We developed a protocol for enzymatic dissociation of fresh mouse lungs based on multiple published techniques [3-5]. Mice were perfused with 10mL 1X Hank's balanced salt solution (HBSS; Corning CellGro, No. 21-023-CV). Lungs were collected and rinsed in chilled 1X PBS. The tissue was cut into small pieces with a razor blade and incubated in 5mg/mL type 2 collagenase (Worthington Biochemical Corporation, No. LS004174) in phenol-red free DMEM for 10min at 37ºC on a benchtop shaker. Enzyme A/Buffer Y mix from the Neural Tissue Dissociation Kit P from Miltenyi Biotec (No. 130-092-628) was added to the collagenase-tissue mixture and incubated for another 15min at 37ºC on a benchtop shaker. Pipetting periodically during this incubation was used to dissociate the tissue. Ice cold PBS/bovine serum albumin (BSA) cell buffer (1X PBS with 0.5% 50mg/mL BSA; ThermoFisher Scientific, No. AM26160) was used to halt the dissociation enzymatic reaction. Cells were passed through a 100μm mesh filter and centrifuged at 300 xg for 5min at 4ºC. Cells were then resuspended in ammonium-chloride-potassium (ACK) buffer for 1min at 20ºC to lyse any remaining red blood cells. Cold PBS/BSA cell buffer was added and cells centrifuged at 300 xg for 10min at 4ºC. Cells were resuspended in cold 1X PBS and debris was removed using cold Debris Removal Solution (Miltenyi Biotech, No. 130-109-398) according to the manufacturer's instructions. Cells were washed again with PBS/BSA cell buffer and centrifuged at 300 xg for 5min at 4ºC. Finally, cells were resuspended in 500μL-1mL PBS/BSA cell buffer and filtered through a 40μm Flowmi cell strainer (Millipore Sigma, No. BAH136800040).

Gene set overlap. The BioVenn web application (https://www.biovenn.nl/) was used to compare gene lists between RNAseq datasets [6]. Input gene lists included the mouse AT2 differentially expressed genes (high metastatic burden vs. low metastatic burden) with $p \leq 0.05$ and human iAT2 differentially expressed genes (co-cultured vs. cultured alone) with adjusted $p \leq 0.05$.

Functional pathway analysis. The NIH DAVID bioinformatics tool (https://davidbioinformatics.nih.gov/) was used to perform functional pathway analysis on RNAseq datasets [7, 8]. Input gene lists included the mouse AT2 differentially expressed genes (high metastatic burden vs. low metastatic burden) with $p \leq 0.05$ and human iAT2 differentially expressed genes (co-cultured vs. cultured alone) with adjusted $p \leq 0.05$. Data includes functional annotation clusters with enrichment scores >2.0 from each gene list that were then compared across all datasets.

Upstream regulator analysis. Ingenuity pathway analysis (IPA) upstream regulator analysis [9] was used to examine predicted upstream regulators of genes identified by NIH DAVID pathway analysis to be included in the secreted/signaling peptide pathway.

Publicly available scRNAseq analysis. The CZ CELLxGENE Discover online tool (https://cellxgene.cziscience.com) was used to explore single cell gene expression in publicly available datasets [10]. The GENE Discover collection currently includes 1338 human and 389 mouse published scRNAseq datasets.

**Statistical analysis.** Differences between two groups were determined by unpaired two-tailed *t*-tests with a Welch's correction when an F-test of equality for variance showed significant differences in variance between groups. Differences between two groups for multiple targets were determined by multiple unpaired two-tailed *t*-tests, with Welch's corrections when appropriate, and the false discovery rate (FDR) controlled using the two-stage step-up (Benjamini, Kreieger, and Yekutieli) method [11]. Differences between multiple groups was determined by ordinary one-way ANOVA with Tukey's multiple comparisons test. Differences between multiple groups for multiple targets were determined by two-way ANOVA with Sidak's multiple comparison test. Linear regressions were used to analyze correlative comparisons with the proportion of variance from a linear relationship presented as $R^2$.
