## Supplementary Figures for "Metastasis-associated wound repair promotes reciprocal activation of the lung epithelium and breast cancer metastases during outgrowth"

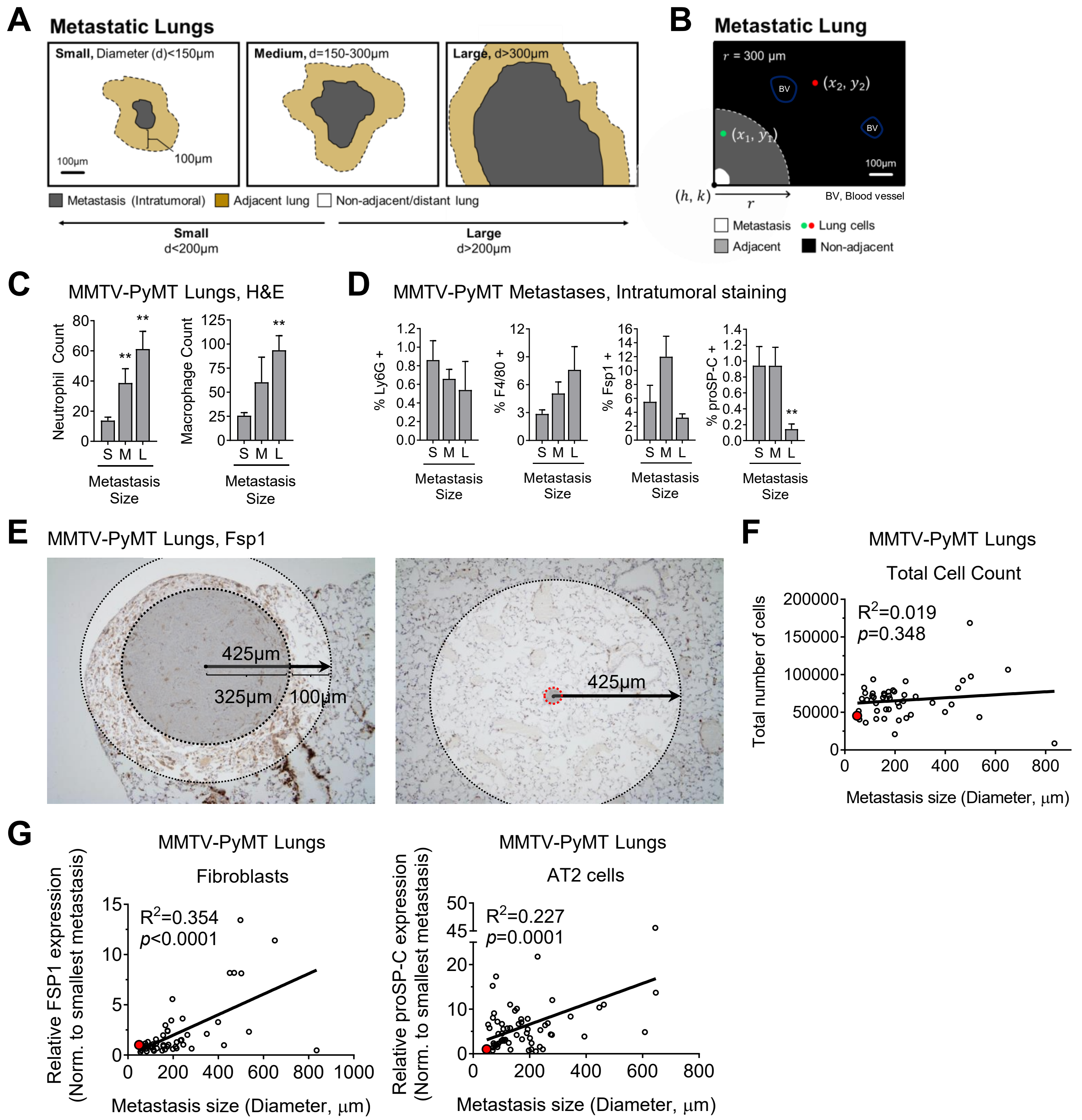


### Supplementary Figure 1.

**Supplementary Figure 1. Histological analysis of lung metastases. A,** Model defining metastasis size designations based on diameter in µm and classifying metastasis adjacent versus non-adjacent/distant tissue in the lung microenvironment for IHC stains. **B,** Model depicting how multispectral immunofluorescence data was quantified and how cells were determined to be metastasis adjacent versus non-adjacent. **C,** MMTV-PyMT metastatic lungs were H&E stained ($n$=20 metastases from 3-4 mice per stain; see Supplementary Table 1 for sample number details). The number of neutrophils and macrophages adjacent to metastases were counted by a veterinary pathologist. Mean ± SEM (unpaired *t*-tests with Welch's correction when appropriate); ** $p$<0.01. S, small; M, medium; L, large metastases. **D,** MMTV-PyMT metastatic lungs were stained for cell-specific markers of lung wound repair ($n$ >50 metastases from 2-7 mice per stain). The percentage of positively stained intratumoral metastatic cells, normalized to the total number of cells, was quantified. Mean ± SEM (unpaired *t*-tests with Welch's correction when appropriate); ** $p$<0.01. S, small; M, medium; L, large metastases. **E,** Images depicting how lung compaction surrounding growing metastases was investigated in MMTV-PyMT metastatic lungs stained for the fibroblast marker Fsp1 and the AT2 marker proSP-C by IHC. The 425µm surrounding each metastasis was determined by measuring the 100µm surrounding the largest lung metastases in our dataset and analyzed for positively stained cells. **F,** The total number of cells surrounding metastases in MMTV-PyMT lungs was quantified in lungs stained for Fsp1. Total number of cells relative to metastasis size for each metastasis was quantified. The red circle represents data for the smallest metastasis within our dataset (linear regression analysis from n>49 metastases in 7 mice). **G,** The number of positively stained cells surrounding metastases in MMTV-PyMT lungs was normalized to the data from the smallest metastasis (red circle) and correlated to metastasis size (linear regression analysis from n>49 metastases in 7 mice).

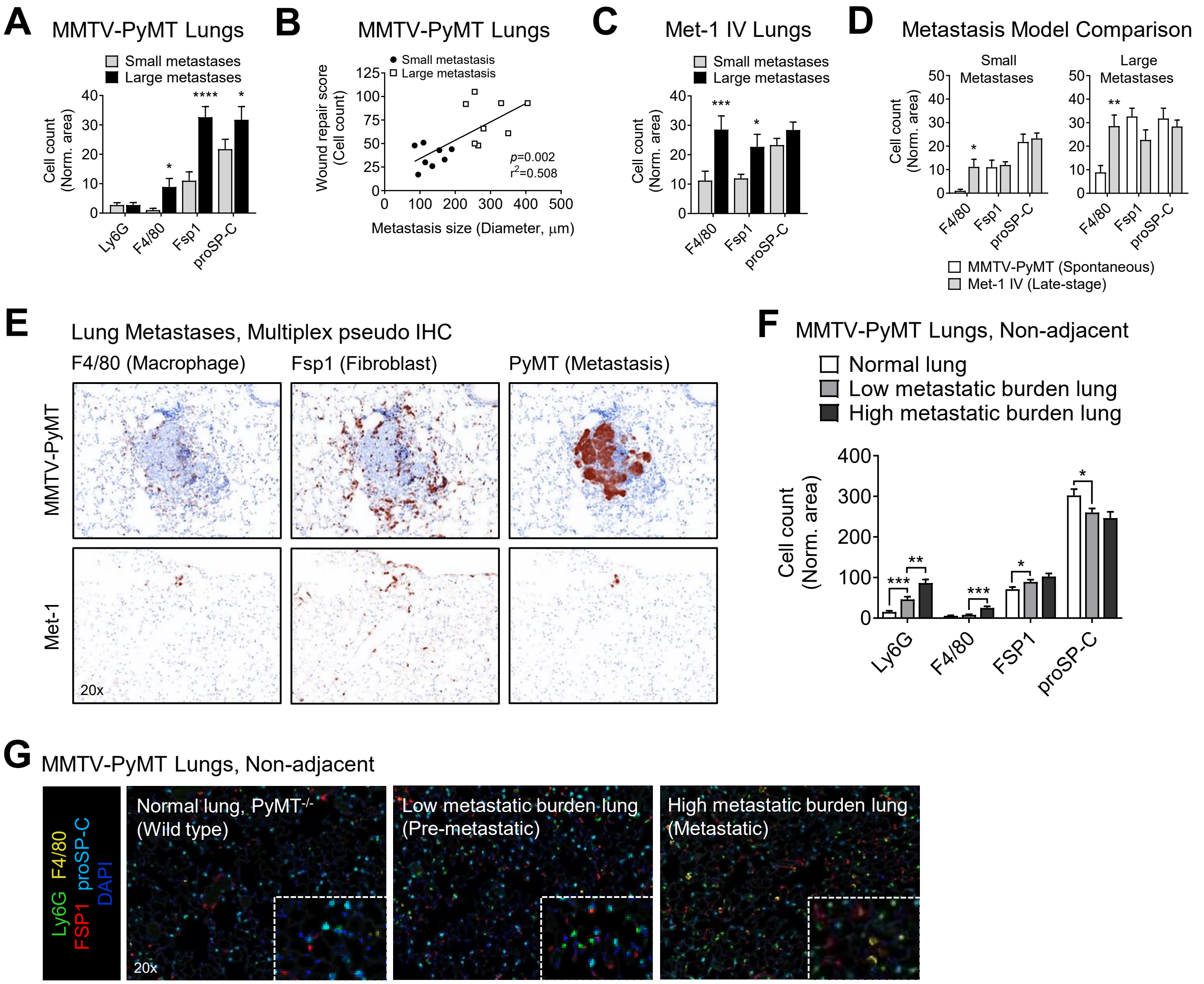


**Supplementary Figure 2. Multispectral immunofluorescent analysis of lung metastases. A,** MMTV-PyMT metastatic lungs were stained for cell-specific markers of lung wound repair (*n*=16 metastases from 2-4 mice per group). The number of positively stained cells, normalized to area, was quantified in the 300µm surrounding metastases. Mean ± SEM (multiple unpaired *t*-tests); * $p \leq 0.05$, **** $p < 0.0001$. **B,** Wound repair scores from MMTV-PyMT lungs relative to metastasis size (linear regression from *n*=16 metastases in 2-4 mice). **C,** Met-1 metastatic lungs were stained for cell-specific markers of lung wound repair (*n*=50 metastases from 5 mice). The number of positively stained cells, normalized to area, was quantified in the 300µm surrounding metastases. Mean ± SEM (multiple unpaired *t*-tests); * $p \leq 0.05$, *** $p < 0.001$. **D,** The number of positively stained cells surrounding metastases, normalized to area, was compared in the spontaneous MMTV-PyMT metastasis model (*n*=16 metastases from 2-4 mice) and the late-stage Met-1 IV metastasis model (*n*=50 metastases from 5 mice). Mean ± SEM (multiple unpaired *t*-tests); * $p \leq 0.05$, ** $p < 0.01$. **E,** Lungs from MMTV-PyMT and Met-1 metastasis models were stained by multispectral immunofluorescence for cell-specific markers of wound repair. Shown are representative pseudo-colored images using a 20X objective. **F-G,** Lungs from wild-type mice (PyMT$^{-/-}$) and MMTV-PyMT mice with low (pre-metastatic) or high metastatic burden were stained for cell-specific markers of wound repair. The number of positively stained cells, normalized to area, was quantified in non-adjacent lung tissue (n=40 regions of interest from 3 mice). Mean ± SEM (multiple unpaired *t*-tests); * $p \leq 0.05$, ** $p < 0.01$, *** $p < 0.001$. Shown are representative images using a 20X objective, inset zoom 2x.

### Supplementary Figure 2.

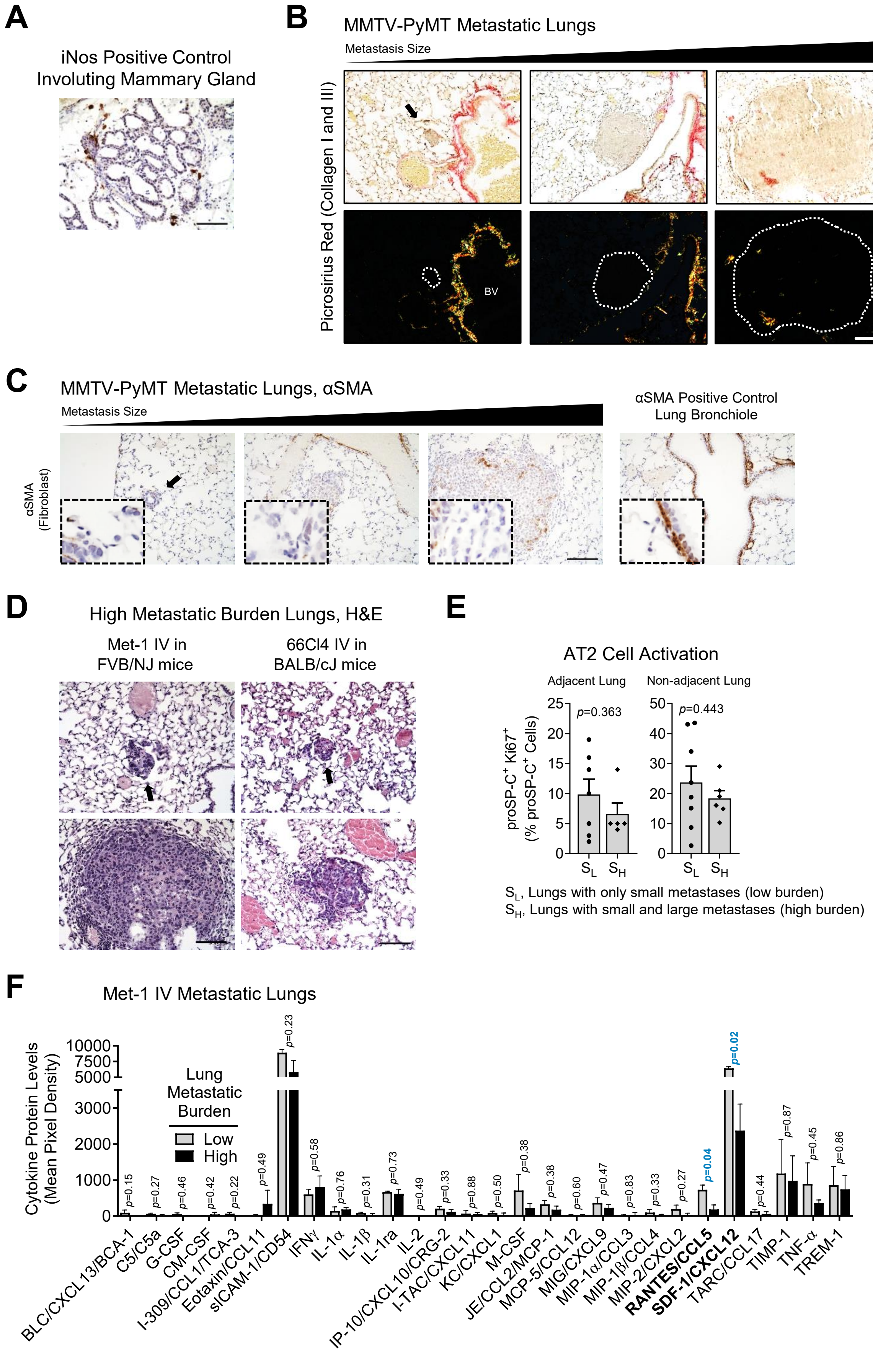


**Supplementary Figure 3.**

**Supplementary Figure 3. Cell activation in metastatic lungs. A,** Representative image of iNOS staining in an involuting mouse mammary gland used as a positive control for immunohistochemical staining; scale bar = 100µm. **B,** Representative images from MMTV-PyMT metastases of different sizes with black arrow indicating small metastases stained with picrosirius red. Metastases are outlined in white in polarized light images. BV, blood vessel; scale bar = 100µm. **C,** MMTV-PyMT metastatic lungs were stained for the fibroblast activation marker αSMA. Shown are representative images from metastases of different sizes with black arrows indicating small metastases; scale bar = 100µm, inset zoom 4x. **D,** Representative images of H&E stained high metastatic burden lungs from late-stage metastasis models using Met-1 cells in FVB/NJ mice and 66Cl4 cells in BALB/cJ mice; scale bar = 100µm. **E,** Multispectral immunofluorescent staining of MMTV-PyMT metastatic lungs for AT2 cell activation. The percentage of proSP-C-positive cells that are Ki67-positive was quantified in the adjacent and non-adjacent lungs surrounding small metastases from low metastatic burden lungs with only small metastases ($S_L$) and high metastatic burden lungs with both small and large metastases ($S_H$) (*n*=14 metastases from 2-3 mice; unpaired *t*-tests), mean $\pm$ SEM. **F,** Cytokine array performed on lungs from mice with a low or high metastatic burden using the late-stage Met-1 metastasis model (*n*=3 mice per group), mean $\pm$ SEM (multiple unpaired *t*-tests with Welch's correction).

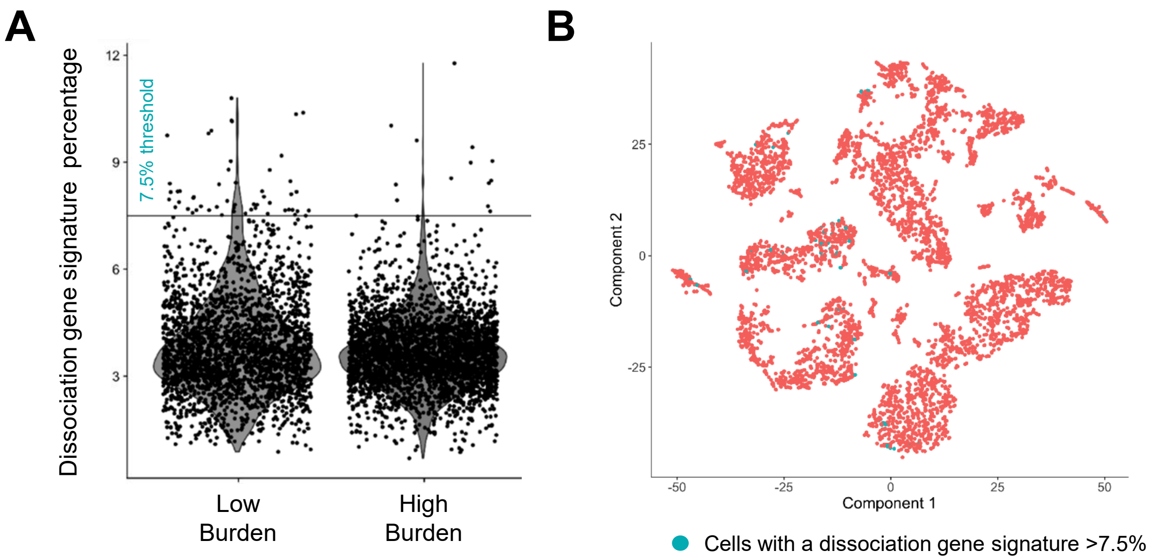


**Supplementary Figure 4. scRNAseq dissociation gene signature analysis.** Lungs from mice with a low or high metastatic burden using the late-stage Met-1 metastasis model were enzymatically dissociated for downstream scRNAseq analysis (n=1 mouse per group). **A,** The dissociation percentage (PMID: 28960196) was calculated for each cell by examining the counts of each dissociation-related gene (138) relative to the total counts of all genes per cell. 65 cells out of 5504 have a dissociation percentage greater than the published threshold of 7.5%. **B,** t-SNE visualization of cells with a dissociation percentage >7.5% (colored blue). Only 2% of cells from a low metastatic burden lung and 0.4% of cells from high metastatic burden lung have a dissociation signature >7.5%.

### Supplementary Figure 4.

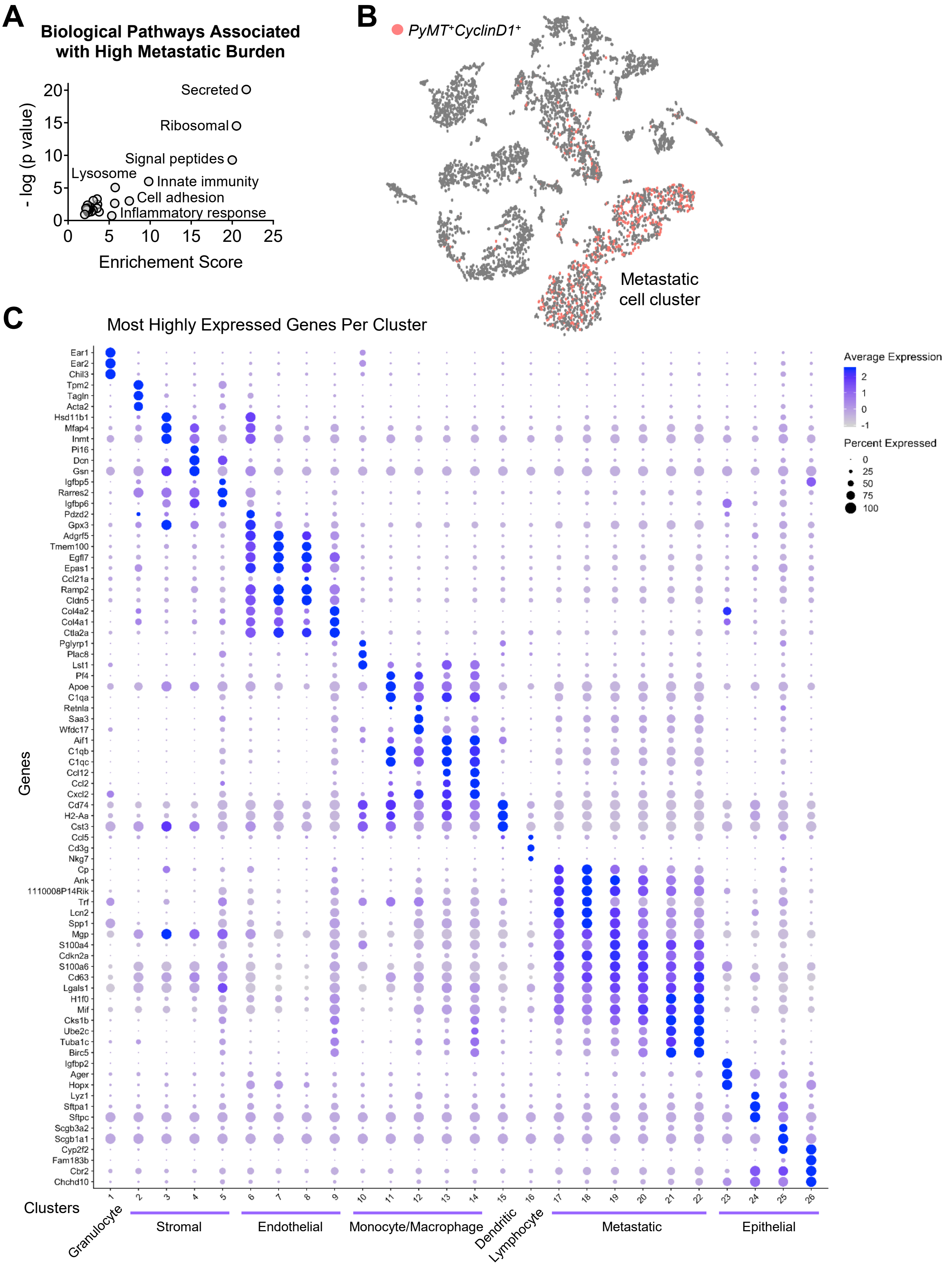


**Supplementary Figure 5.**

**Supplementary Figure 5. scRNAseq analysis of metastatic outgrowth in lungs.** Lungs from mice with a low or high metastatic burden using the late-stage Met-1 metastasis model were transcriptionally evaluated using scRNAseq (n=1 mouse per group). **A,** A bulk analysis was performed on the combined data from all cells in lungs with a low metastatic burden compared to cells from lungs with a high metastatic burden. Functional biological pathways associated with metastatic outgrowth were identified using differentially expressed genes. Only pathway clusters with an enrichment score >2 are included. **B,** Identification of the mammary carcinoma metastatic cell population. t-SNE visualization of lung cells clustered by gene expression and colored by co-expression of *PyMT* and *CyclinD1*. **C,** The most highly expressed genes per cell cluster. Average expression defined as average log fold change between one cluster and all other clusters. Percent expression defined as the percentage of cells within each cluster that express each gene.

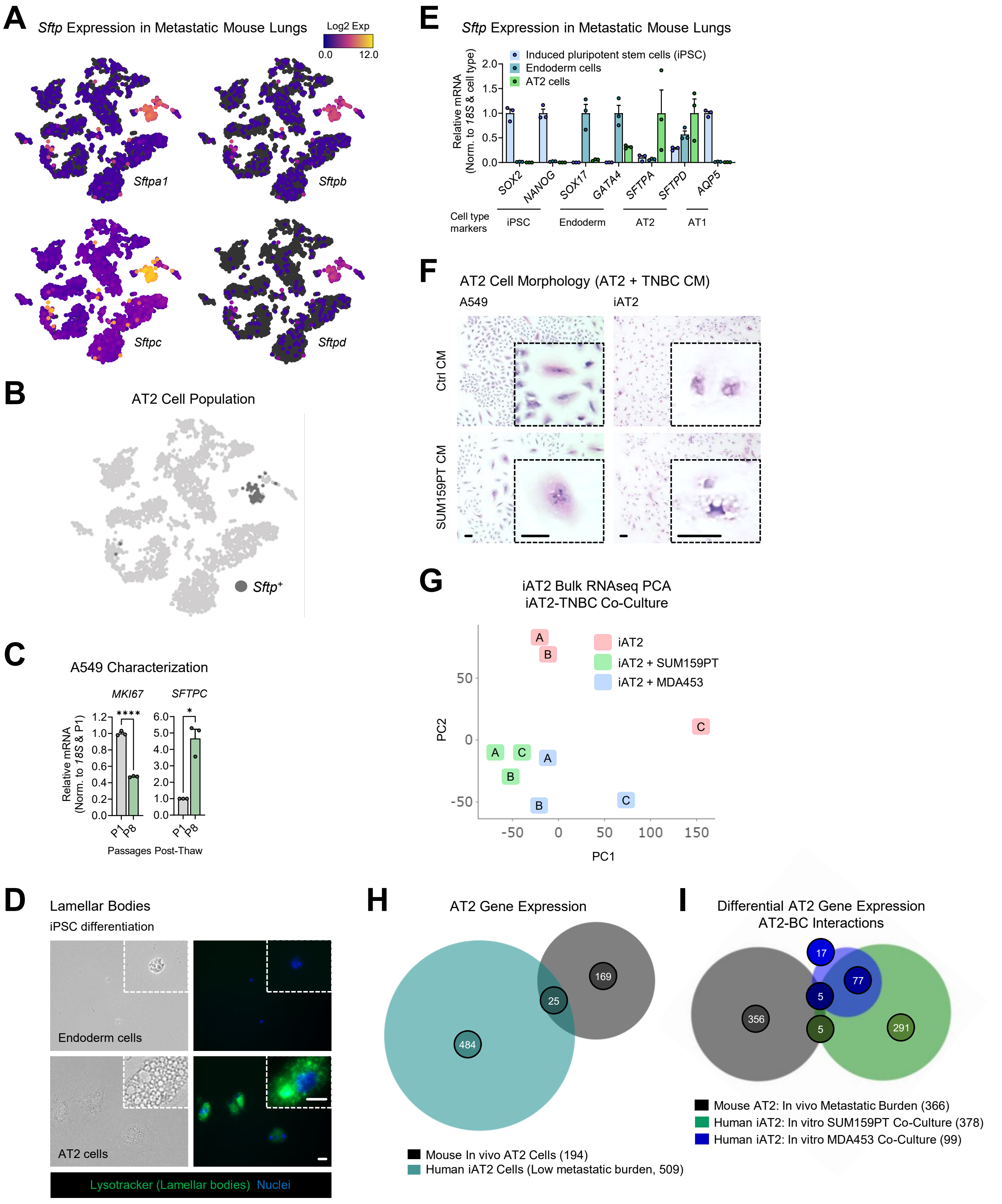


### Supplementary Figure 6

**Supplementary Figure 6. Lung AT2 cells. A,** Lungs from mice with a low or high metastatic burden using the late-stage Met-1 metastasis model were transcriptionally evaluated using scRNAseq (n=1 mouse per group). t-SNE visualization of surfactant protein (*Sftp*) isoform gene expression. **B,** Identification of the AT2 cell population. t-SNE visualization of combined *Sftpa-d* gene expression (*Sftp*$^{+}$) with a log2 max expression >12. **C,** Gene expression by qPCR of A549 cells following long-term culture (passage 8, P8). Data was normalized to *18S* and early passage A549 cells (passage, P1). Mean $\pm$ SEM (unpaired *t*-tests with Welch's correction when appropriate); * $p \leq 0.05$, **** $p < 0.0001$. **D,** iAT2 lamellar bodies were imaged using lysotracker staining following induced pluripotent stem cell (iPSC) differentiation through the endoderm stage to AT2 cells; scale bar = 10μm, inset zoom 3x. **E,** Gene expression by qPCR in iPSC, endoderm, and iAT2 cells for cell type specific markers of differentiation. Data was normalized to *18S* and cell type for each marker; mean $\pm$ SEM. **F,** AT2 cells were cultured in 50% conditioned media (CM) from SUM159PT cells or control (Ctrl) media for 3 days. AT2 cell morphology was then examined by H&E. Shown are representative images of cell size and shape; scale bar = 10μm, inset zoom 4x for A549 cells and 6x for iAT2 cells. **G,** Bulk RNAseq was performed on RNA collected from iAT2 cells co-cultured with SUM159PT or MDA453 cells for 5 days. Control iAT2 cells were cultured alone. This principal component analysis (PCA) illustrates the relationship between gene expression data from sample/replicates. **H,** Venn diagram comparing genes expressed in AT2 cells in mouse lungs (data from mouse lungs with a low metastatic burden) versus genes expressed in human iAT2 cells cultured alone. **i,** Venn diagram comparing AT2 genes associated with BC cell interactions using differentially expressed genes from mouse AT2 cells in metastatic lungs and human iAT2 cells co-culture with TNBC cells.

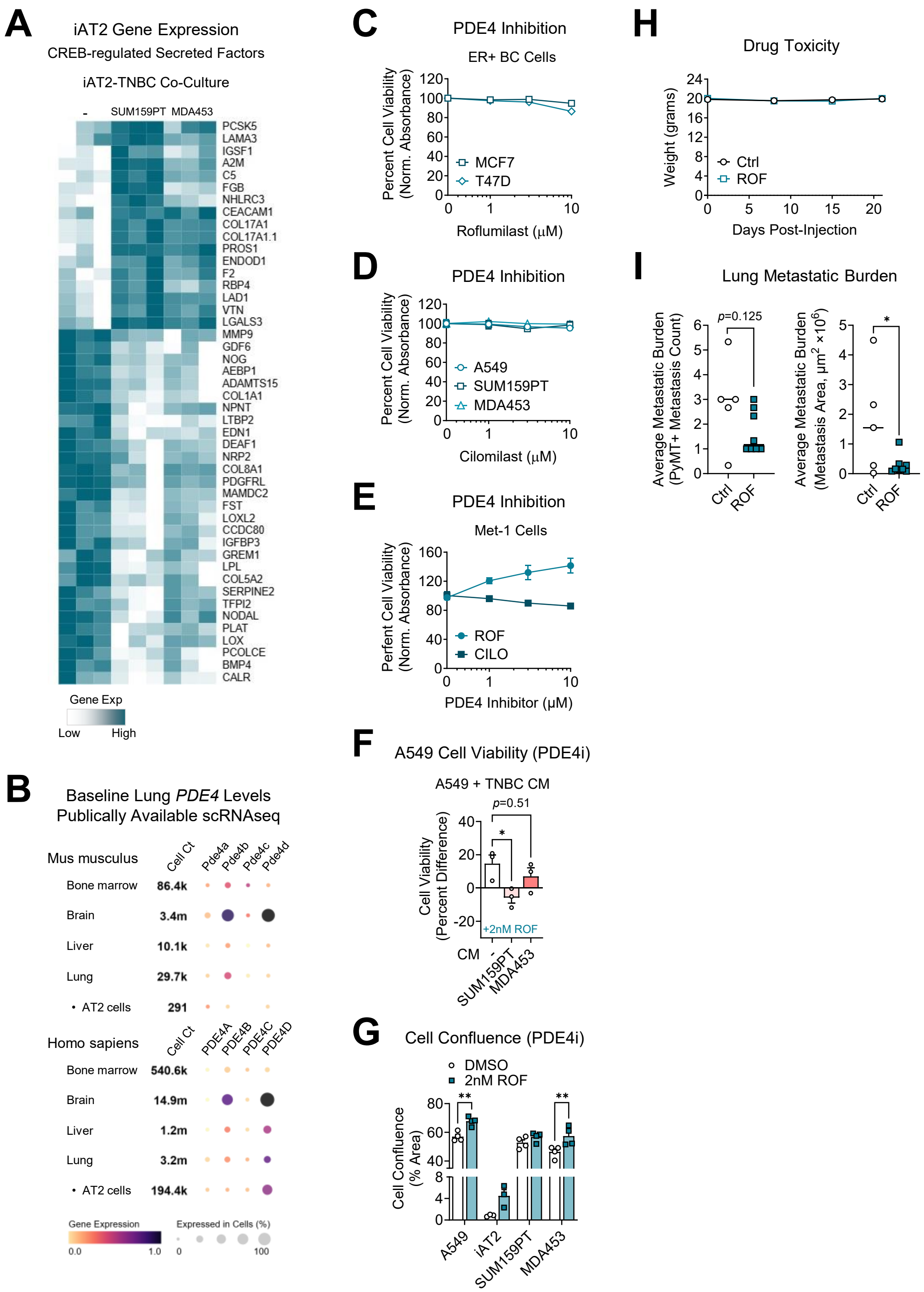


### Supplementary Figure 7

**Supplementary Figure 7. PDE4 levels and inhibition. A,** Heatmap of CREB-regulated secreted factor gene expression in iAT2 cells cultured alone (-) or co-cultured with SUM159PT or MDA453 cells for 5 days. **B,** Publicly available scRNAseq data was queried using the CZ CELLxGENE Discover data platform for *PDE4* isoform baseline levels in mouse and human tissue and cells (PMID: 390607691). **C,** Percent cell viability was measured by crystal violet assay after 3 day treatment with vehicle DMSO (0) or increasing concentrations of the PDE4 inhibitor roflumilast (ROF). Data was normalized to mean absorbance of DMSO-treated cells; mean ± SEM. **D,** Percent cell viability was measured by crystal violet assay after a 3 day treatment with vehicle DMSO (0) or increasing concentrations of the PDE4 inhibitor cilomilast (CILO). Data was normalized to the mean absorbance of DMSO-treated cells; mean ± SEM. **E,** Percent cell viability was measured in Met-1 mouse carcinoma cells by crystal violet assay after a 3 day treatment with vehicle DMSO (0) or increasing concentrations of ROF or CILO. Data was normalized to the mean absorbance of DMSO-treated cells; mean ± SEM. **F,** A549 cell viability was measured by crystal violet assay following culture with conditioned media (CM) from TNBC cells and treatment with 2nM roflumilast (ROF) for 5 days. The percent difference in cell viability was calculated. Mean ± SEM (one-way ANOVA with Tukey's multiple comparison test); * $p \leq 0.05$. **G,** Cell confluence was examined by crystal violet assay in cells treated with DMSO or 2nM ROF for 5-7 days, depending on the cell line. Mean ± SEM (two-way ANOVA with Sidak's multiple comparison test); ** $p < 0.01$. **H,** Met-1 cells were injected IV into the tail veins of female mice and orally administered 5mg/kg ROF starting 3 days post-injection ($n$=5-8 mice per group). Mice were treated daily for 3 weeks and drug toxicity was tracked through weekly measurement of mouse weight; mean ± SEM. **i,** Metastatic lung tissue was collected from ROF treated mice and stained for the Met-1 mammary-specific marker PyMT. Average metastatic burden was quantified in serial sections as the number of PyMT+ metastases or the size of PyMT+ metastases as measure by area in $um^2$. The average of three serial sections was calculated per mouse. Mean ± SEM ($n$=5-8 mice per group, unpaired $t$-tests); * $p \leq 0.05$.
